# A symmetric toggle switch explains the onset of random X inactivation in different mammals

**DOI:** 10.1101/204909

**Authors:** Verena Mutzel, Ikuhiro Okamoto, Ilona Dunkel, Mitinori Saitou, Luca Giorgetti, Edith Heard, Edda G. Schulz

## Abstract

Gene-regulatory networks control establishment and maintenance of alternative gene expression states during development. A particular challenge is the acquisition of opposing states by two copies of the same gene, as it is the case in mammals for *Xist* at the onset of random X-chromosome inactivation (XCI). The regulatory principles that lead to stable mono-allelic expression of *Xist* remain unknown. Here, we uncovered the minimal *Xist* regulatory network, by combining mathematical modeling and experimental validation of central model predictions. We identified a symmetric toggle switch as the basis for random mono-allelic *Xist* up-regulation, which reproduces data from several mutant, aneuploid and polyploid murine cell lines with various *Xist* expression patterns. Moreover, this toggle switch explains the diversity of strategies employed by different species at the onset of XCI. In addition to providing a unifying conceptual framework to explore X-chromosome inactivation across mammals, our study sets the stage for identifying the molecular mechanisms required to initiate random XCI.

## Introduction

During developmental cell fate decisions, each cell must choose between alternative transcriptional states that must then be stably maintained. One of such decision-making processes occurs at the onset of random X-chromosome inactivation (XCI), where half of the cells in female embryos will silence the maternal (Xm) and the other half the paternal X chromosome (Xp). XCI is initiated during early embryogenesis by mono-allelic up-regulation of the long non-coding RNA (lncRNA) *Xist* from either the Xp or the Xm, which then induces chromosome-wide gene silencing *in cis*. *Xist* recruits several repressive histone modifications including H3K27me3 to the inactive X, eventually resulting in complete heterochromatinization of the entire chromosome. In this way mammals ensure dosage compensation for X-linked genes between the sexes^1^.

While *Xist* appears to control XCI in all eutherian mammals, marked differences have been found in the way it is regulated^2^. Human and rabbit embryos pass through an initial stage where *Xist* is expressed from both X chromosomes^3^. By contrast, *Xist* up-regulation in mouse is generally thought to be strictly mono-allelic^4,5^. While the bi-allelic phase is very transient in rabbits, it extends over several days in human embryos, where it does not induce gene silencing, but only dampening of gene expression^3,6^. The regulators of *Xist* also appear to be rather poorly conserved across species^2^. *Tsix, Xist’s* repressive antisense transcription unit, plays an important role in mice, but might not be functional in other mammals^2,7,8^, while humans have evolved another lncRNA that antagonizes *Xist*, called *XACT*^9^. Therefore, it has been suggested that different species employ diverse strategies for the initial establishment of XCI during embryogenesis^2^.

To establish the female-specific mono-allelic expression pattern of *Xist*, a cell must (i) assess the number of X chromosomes that are present, (ii) choose one of them for *Xist* up-regulation and (iii) stabilize the two opposing states of the inactive X (Xi), which expresses *Xist*, and the active X (Xa) where *Xist* is silent^1^. The underlying regulatory network appears to integrate information on X-chromosomal dosage, since *Xist* is up-regulated in cells with two or more X chromosomes, but not in male or XO cells with only a single X^10^. Interestingly, diploid cells with four X chromosomes (X tetrasomy) inactivate three X’s, while tetraploid cells that also contain four X chromosomes only inactivate two of them, suggesting that autosomal ploidy modulates the onset of XCI as well^10,11^.

Since the two *Xist* loci in female cells adopt opposing expression states, major regulatory events must occur *in cis* on the allele level. Central regulators of *Xist* are thus expected to be encoded by X-linked genes, which can mediate *cis*-regulation and transmit X-dosage information. Several *cis*-acting lncRNA loci have been identified, functioning either as *Xist* repressors, such as *Tsix* and potentially *Linx* or as *Xist* activators, such as *Ftx* and *Jpx*^7,12–14^. X-dosage sensing has been suggested to be mediated by a *trans*-acting X-linked *Xist* activator, which would be present in a double dose in female compared to male cells and could thus confer female-specificity of XCI^1^. Silencing of XA upon mono-allelic *Xist* up-regulation would then prevent *Xist* expression from the other allele, thus forming a trans-acting negative feedback loop^15,16^. Two *trans*-acting *Xist* activators have been proposed so far, the Rlim/Rnf12 protein, which is silenced by *Xist* and the lncRNA *Jpx*, which escapes XCI^13,15,17^.

A series of regulators involved in the initiation of XCI have thus been discovered over the years, but each one has mostly been studied in isolation. Their relative contributions and functional interplay, as well as the underlying regulatory principles remain poorly understood. Here, to rigorously identify the interactions required for the initiation of random XCI we use a novel approach, where we employ mathematical modeling and simulations to compare alternative network architectures and test model predictions experimentally. We show that two different *Xist* regulators, a *cis*-acting repressor and a *trans*-acting activator, are sufficient and must cooperate to ensure female-specific mono-allelic up-regulation of *Xist*. They form an extended symmetric toggle switch, which can explain *Xist* expression in aneuploid and polyploid cells as well as the seemingly diverse *Xist* patterns in different species. Moreover, we show that in mice, the predicted cis-acting repressor could be *Xist’s* antisense transcription unit *Tsix*. Our systems biology approach has thus identified the regulatory principles governing the onset of XCI and provides a unifying framework of *Xist* regulation across species.

## Results

### Identifying a core network that can maintain mono-allelic *Xist* expression

To investigate the regulatory principles governing mono-allelic and female-specific *Xist* expression, we systematically screened alternative architectures of the underlying regulatory network. To this end, potential X-linked *Xist* regulators were classified in 8 different categories depending on whether they activate (A) or repress (R) *Xist*, whether they act *in cis* (c) on the same chromosome or *in trans* (t) on both chromosomes and whether they are silenced during XCI or escape (e) (Fig. 1A and Table 1). For each regulator type, we used ordinary differential equations (ODE) to build a mathematical model that describes a cell with two X chromosomes containing *Xist* and the respective regulator (for details see supplemental model description).

To understand which networks are able to maintain mono-allelic *Xist* expression, each model was simulated starting from an XaXi state (simulation 1, Fig. 1B), where *Xist* is highly expressed from the Xi and not expressed from the Xa. The simulation was performed for at least 1000 randomly chosen parameter sets, combining different values for transcription rates and activation/repression strengths, to test whether the network could in principle reproduce the experimental behavior, at least for certain parameter values (for details see supplemental model description). Only the network with a *cis*-acting *Xist* repressor (cXR) was able to maintain mono-allelic *Xist* expression (in 32% of parameter combinations, Fig. 1B). We further tested another 28 models, each combining two regulator types instead of one. In this way, 7 additional, more complex models were identified that could stabilize the mono-allelic state (for details see supplemental model description). Since all of them contained cXR, this appears to be only factor strictly required to maintain the XaXi state.

**Table 1.** X-linked *Xist* regulators. All known X-linked *Xist* regulators are classified according to whether they activate or repress *Xist* (XA/XR), whether they act *in cis* or *in trans* (c/t) and whether they escape XCI (e).

| X-linked regulators |  | Putative members |
| --- | --- | --- |
| cXA | *cis*-acting X-linked activator | - |
| ecXA | escaping *cis*-acting X-linked activator | Ftx<sup>14</sup>, Jpx<sup>13,17</sup> |
| tXA | *trans*-acting X-linked activator | Rnf12<sup>15,18,19</sup> |
| etXA | escaping *trans*-acting X-linked activator | Jpx<sup>13,17</sup> |
| cXR | *cis*-acting X-linked repressor | Tsix<sup>7</sup>, Linx<sup>12</sup> |
| ecXR | escaping *cis*-acting X-linked repressor | - |
| tXR | *trans*-acting X-linked repressor | - |
| etXR | escaping *trans*-acting X-linked repressor | - |

Apart from maintaining the mono-allelic expression state in female cells, the *Xist* regulatory network must ensure that *Xist* is not expressed in male cells and that erroneous bi-allelic *Xist* up-regulation from both chromosomes in females is prevented. To test which network would fulfill these criteria we performed three additional simulations, where we tested whether male cells with a single X chromosome would maintain the Xa state (simulation 2) and whether the erroneous XiXi and Xi states in female (simulation 3) and male cells (simulation 4), respectively, would be unstable. These simulations were performed for all 8 models that maintained the mono-allelic state in simulation 1, which contained cXR either alone or in combination with another regulator type (Fig. 1C). While all tested networks maintained the Xa state in simulation a *trans*-acting *Xist* activator (tXA) was required to prevent erroneous *Xist* expression in simulations 3 and 4 (Fig. 1C). To ensure female-specificity of *Xist* up-regulation, it is irrelevant whether tXA is silenced by *Xist* (simulation 4); however to prevent bi-allelic expression in female cells it must be subject to XCI (simulation 3). Through comprehensive screening of 36 alternative network architectures we have thus identified a single minimal network that can ensure the correct *Xist* expression pattern, where a *trans*-activating activator (tXA) that is silenced by *Xist* must cooperate with a *cis*-acting *Xist* repressor (cXR, Fig. 2A). Although the *trans*-activator hypothesis has been previously proposed^11^, our analysis reveals that without a *cis*-acting repressor this would not be enough to ensure correct XCI.

**Figure 1:**
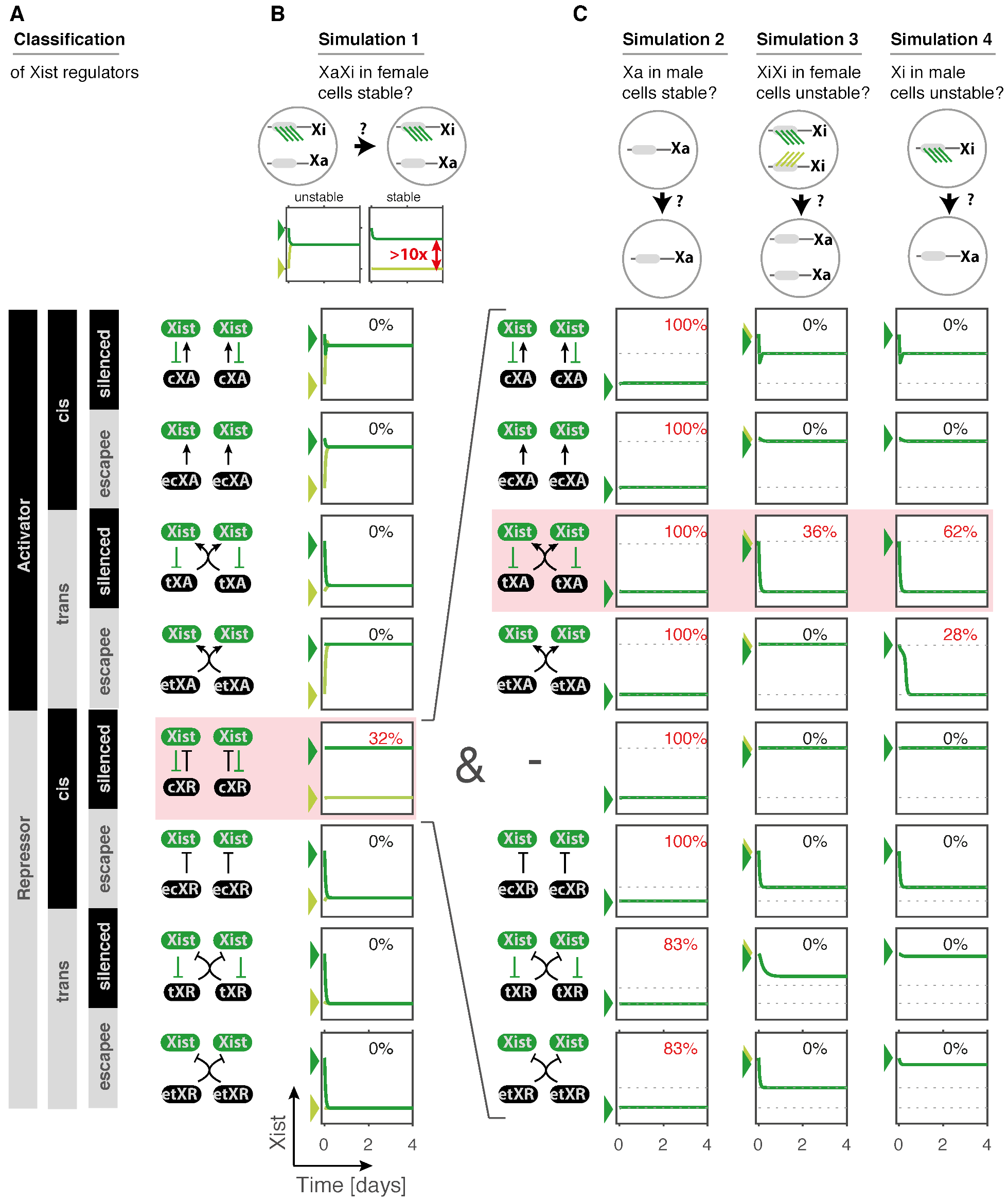
Comparison of alternative model structures. **(A)** Schematic depiction of the classification of X-linked *Xist* regulators, depending on whether they act as activators (XA) or repressors (XR), whether they act *in cis* (c) or *in trans* (t) and whether they are silenced during XCI or escape (e). The networks that are formed by *Xist* and each regulator are shown schematically (right). **(B)** To test which network could stabilize mono-allelic *Xist* expression, they were translated into mathematical models (ODE), describing two X chromosomes, each carrying *Xist* and the respective regulator. Each model was simulated with >1000 randomly chosen parameter sets initiating from an XaXi state (scheme, top), where *Xist* is expressed from one chromosome (dark green), and not from the other (light green). One example simulation is shown for each network and the percentage of tested parameter sets, where the XaXi state was stably maintained is indicated (in red if >0%). **(C)** The *cis*-acting repressor (cXR) that could maintain the XaXi state in (B) was combined with all other regulator classes to build 7 more complex models. For all parameter sets that could maintain the XaXi state three additional simulations were performed to test whether the Xist^OFF^ state (Xa) was maintained in male cells with a single X chromosome (simulation 2), whether bi-allelic *Xist* expression (XiXi) would be unstable in female cells (simulation 3) and whether *Xist* expression from the single X (Xi) in male cells would be unstable (simulation 4). One example simulation is shown for each model and the percentage of parameter sets that fulfill these criteria are indicated. Dotted lines indicate the Xa and Xi state from the simulation in (B) and arrow heads denote the initial conditions. Shaded boxes indicate the model that can reproduce the experimental observations.

### The cXR-tXA model can reproduce initiation of XCI in diploid, polyploid and polysomic cells

We then asked whether the identified network (Fig. 2A) could also recapitulate monoallelic *Xist* up-regulation at the onset of X inactivation. The transition from the pre-XCI XaXa state to the post-XCI XaXi state, where either the Xm or Xp are randomly inactivated, cannot be simulated in a deterministic framework such as the ODE systems used above. We thus developed a stochastic model of the cXR-tXA network, which simulates individual cells and takes random fluctuations into account (details given in supplemental model description). When initiating the simulation from an XaXa state (Fig. 2B) a subset of parameter sets was able to reproduce robust mono-allelic upregulation of *Xist* (example simulations in Fig. 2C+D, detailed analysis of the parameter requirements in supplemental model description). Importantly, individual cells randomly up-regulate *Xist* from either one or the other X chromosome (Fig. 2C, light and dark green).

To understand how the cXR-tXA model ensures mono-allelic *Xist* up-regulation, we analyzed the *Xist* expression states that the system assumes at the single-allele level (Fig. 2E, top) as well as at the cell level (bottom). In the presence of a single tXA dose as in post-XCI cells (XaXi) when one copy of tXA is silenced, an allele can assume either a low or a high expression state of *Xist* corresponding to the Xa and Xi, respectively (Fig. 2E, top). In undifferentiated pre-XCI cells (XaXa), tXA is present in a double dose and Xist is repressed by pluripotency factors. At the onset of differentiation, which is the stage described by our model, this repression is relieved and the only stable state that exists is the Xist-high state, thus leading to *Xist* up-regulation (XaXa, Fig. 2E). By contrast, upon complete silencing of tXA in the XiXi state, only the low *Xist* state is present, such that *Xist* expression will not be sustained (XiXi, Fig. 2E). As a consequence, globally only the two mono-allelic states are stable, where either the maternal or the paternal X chromosome is inactivated (Fig. 2E, bottom). Single-allele and consequently also double-allele bistability requires regulation of *Xist* by cXR and tXA. In the absence of cXR only a single state remains (Fig. 2F), whereas in the absence of dynamic regulation by tXA additional global states appear, because coordination of the two *Xist* loci is lost and both the XaXa and the XiXi states become stable (Fig. 2G). In conclusion, this bistable behavior is generated by mutual repression of *Xist* and cXR, which form a *cis*-acting double-negative (=positive) feedback loop. tXA, which mediates a second, *trans*-acting feedback loop ensures female-specific and mutually exclusive expression of the two *Xist* alleles.

**Figure 2:**
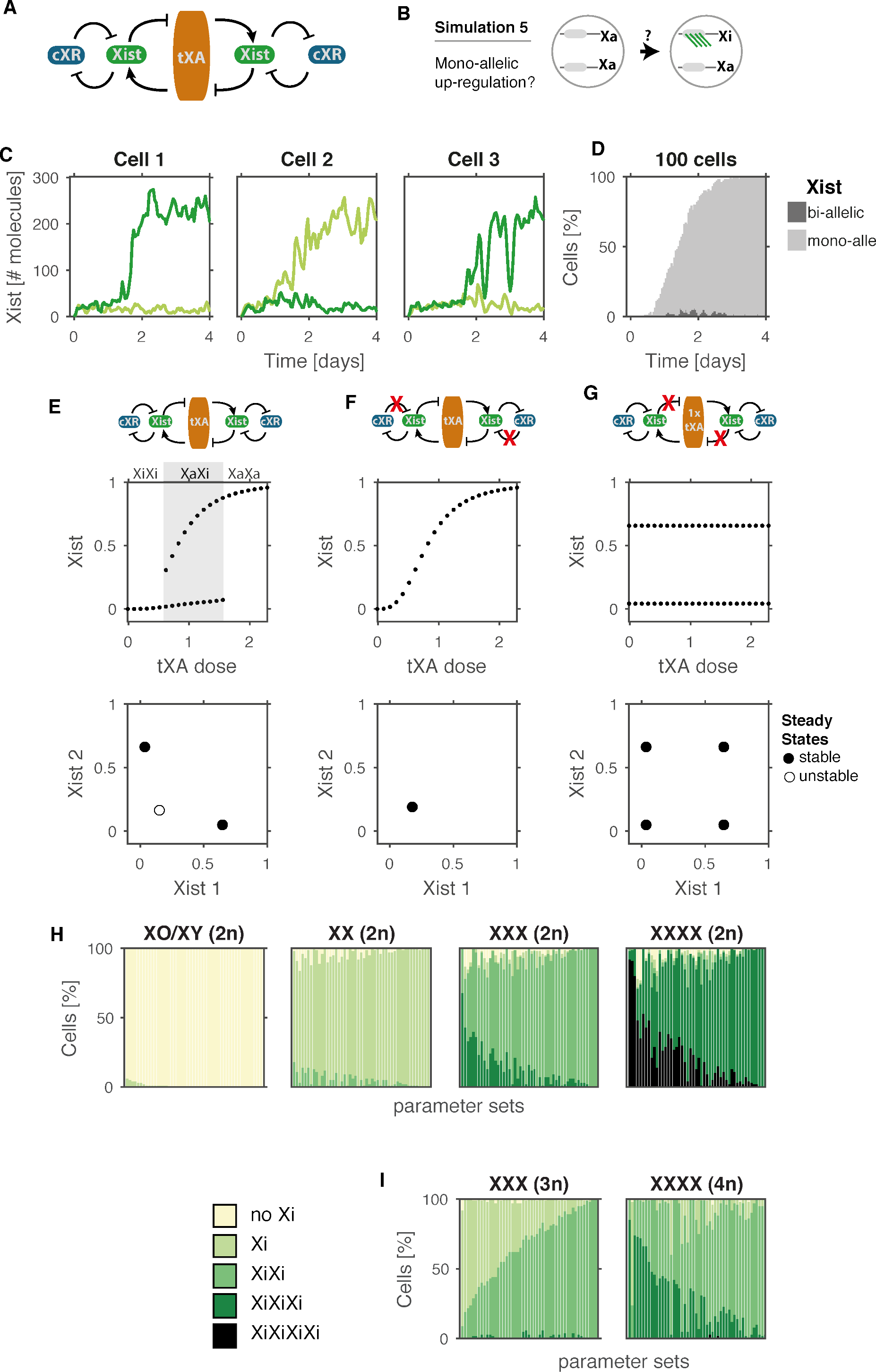
The cXR-tXA model can recapitulate *Xist* patterns in male, female aneuploid and polyploid cell lines. **(A-B)** Schematic representation of the cXR-tXA model (A) and of the stochastic simulation (B) shown in (C-D), which initiates from the XaXa state found in undifferentiated cells. **(C-D)** Simulation of *Xist* up-regulation for one example parameter set, showing three individual cells (C) and a population of 100 cells (D). Light and dark green in (C) represent *Xist* levels expressed from the two X-chromosomes, light and dark grey in (D) represent mono- and bi-allelic *Xist* expression, as indicated. **(E-G)** Steady state *Xist* levels simulated deterministically (as in Fig. 1B) either for the full cXR-tXA model (E) or in the absence of either cXR-mediated repression (F) or tXA mediated activation (G). Local steady states on the allele level for different tXA doses (top) and global steady states (bottom) are shown. Shaded area in (E) indicates the bistable regime for a single tXA dose corresponding to the mono-allelic XaXi state. Filled and open circles indicate stable and unstable steady states, respectively. In (G) tXA was assumed to be present at a constant single tXA (1x tXA). **(H)** Simulations of diploid cells with either one (left, male), two (middle left, female), three (middle right, X-trisomy) or four X chromosomes (right, X-tetrasomy). Stacked bar graphs show the classification of *Xist* patterns in simulations with 50 randomly chosen parameter sets that can generate robust monoallelic *Xist* up-regulation in female diploid cells. **(I)** Simulations of triploid (left) and tetraploid cells (right) assuming that tXA is repressed by autosomal factors in a dose-dependent manner. Simulation of the alternative assumption where tXA is diluted due to increased nuclear volume in polyploid cells is shown in supplementary figure S1 and details on the simulations are given in the supplement.

To further validate the cXR-tXA model, we tested whether it could reproduce the phenotype observed for different X aneuploidies. It is known that in diploid cells (2n), all X chromosomes except one will be ultimately inactivated^10^. If we assume that each additional X chromosome produces an additional dose of tXA (details in supplemental model description), most parameter sets that can reproduce mono-allelic *Xist* up-regulation in diploid female cells correctly predict no *Xist* expression in male and XO cells and bi- and tri-allelic expression in X-chromosome trisomies and tetrasomies, respectively (Fig. 2H). While diploid (2n) cells with four X chromosomes inactivate three of them, tetraploid (4n) cells that also have 4 X’s only inactivate two of them^11^. Similarly, X-trisomic diploid cells inactivate two X chromosomes, while triploid cells are a mixture of cells with one and two Xi^11,20^. To simulate triploid and tetraploid cells, we assumed either that tXA is repressed by autosomal transcription factors or that tXA is diluted due to an increased nuclear volume in polyploid cells, which is two-fold higher in tetraploid compared to diploid cells^21^. In both cases, most parameter sets could correctly predict a mixture of cells with one and two inactive X chromosomes in triploid cells and a majority of cells with two Xi in tetraploid cells (Fig. 2I + S1). Taken together, we have shown that parameter sets that can reproduce mono-allelic *Xist* up-regulation in diploid female cells can also correctly reproduce the different XCI patterns in X tri- and tetrasomies and in tri- and tetraploidies.

### The cXR-tXA model can explain *Xist* patterns in different species

The regulation of *Xist* has been suggested to be rather diverse among species because bi-allelic expression is observed frequently in human and rabbit embryos, but rarely in mice^2^. The single network architecture we have identified can reproduce very different degrees of transient bi-allelic expression (Fig. 3A), depending on the relative time scales of tXA silencing (sil_tXA_) and *Xist* up-regulation (Switch-ON, defined as average time before Xist reaches 20% of the Xist^high^ state). If *Xist* up-regulation is slow (high Switch-ON time), it will normally occur one allele at a time. Subsequent silencing will shift the system to the bistable regime (Fig. 2E) and thereby lock in the mono-allelic state before *Xist* up-regulation from the other X chromosome occurs. This results in a low frequency of bi-allelically expressing cells as observed in mice (Fig. 3A, yellow example). If *Xist* up-regulation is rapid and silencing is slow (long silencing delay siltXA), *Xist* will initially be expressed from two alleles as observed in rabbit embryos (Fig. 3A, purple example). In this scenario the choice of the inactive X can subsequently occur through mono-allelic silencing of tXA and cXR. Alternatively, silencing of both alleles might reverse *Xist* up-regulation completely as *Xist* expression is unstable if both tXA alleles are silenced (see Fig. 2E) such that the cell can undertake a second attempt to reach the mono-allelic state.

**Figure 3:**
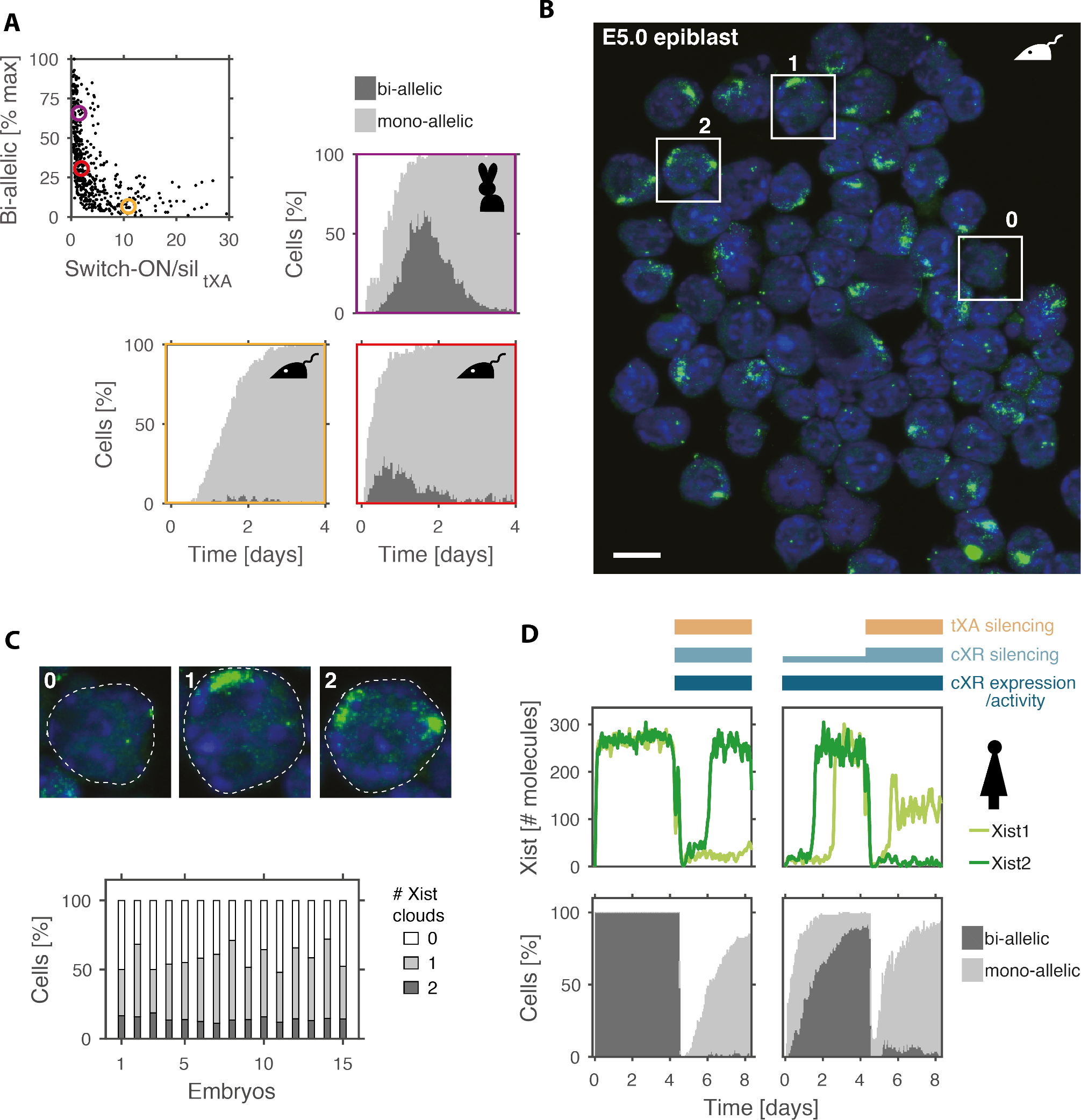
The cXR-tXA model reproduces transient bi-allelic expression in different species. **(A)** For all parameter sets that reproduced mono-allelic *Xist* up-regulation in the cXR-tXA model, the maximal fraction of cells with bi-allelic *Xist* expression observed during the simulation is shown as a function of the the ratio of switch-ON time (first time point, when *Xist* levels reach 20% of the high steady state) and tXA silencing delay (sil_tXA_). Three example simulations with low (yellow), medium (red) and high levels (purple) of bi-allelic expression (dark grey) are shown, thus reproducing observations in mouse and rabbit embryos as indicated. **(B-C)** *Xist/Tsix* RNA FISH (green) and nuclear staining (blue) of female mouse epiblast cells at E5.0 of embryogenesis. (C) Example cells with 0, 1 and 2 *Xist* clouds marked in (B) are enlarged (top) and the number of cells in each category is given across 15 female embryos (bottom). Scale bar (B) 10 μm, dashed white lines (C) indicate the outlines of the nuclei. **(D)** Simulation of bi-allelic expression upon reduced *Xist*-mediated silencing as observed in human embryos, assuming that in the first 4 days of the simulation either silencing and cXR expression/activity is absent (left) or that cXR is silenced partially, while tXA is unaffected by *Xist* (right), as indicated. Simulations of an individual cell (top) and a population of 100 cells (bottom) for one example parameter set are shown. A summary of all parameter sets is given in supplemental figure S2.

Although the same regulatory network can generate different degrees of transient biallelic *Xist* up-regulation, a very low frequency of bi-allelic expression as observed previously in mice (<5%) would require the switch-ON time of *Xist* to be at least 5 to 10 times slower than tXA silencing (Fig. 3A). If we consider that even for rapidly silenced genes the silencing delay is in the range of hours^22,23^, the switch-ON time should be longer than 10h. Such slow dynamics of *Xist* up-regulation, however, are difficult to reconcile with the fact that the transition to the mono-allelic state occurs within 24h between E4.5 and E5.5 of mouse development^4,5,24^. We therefore hypothesized that also in mouse embryos a fraction of cells should up-regulate *Xist* bi-allelically and we set out to study the *Xist* expression pattern *in vivo* in the E5.0 epiblast where random XCI is first initiated (Fig. 3B). RNA FISH analysis revealed that 15-20% of epiblast cells in each embryo exhibited two *Xist* clouds (Fig. 3C). From these data and in agreement with a recent study^25^, we conclude that transient bi-allelic *Xist* expression also occurs during mouse development, but to a lesser extent than in rabbits or humans.

In contrast to rabbits, bi-allelic *Xist* expression in human embryos persists over several days and does not induce gene silencing, but only dampening of gene expression^3,6^. It is unknown how the subsequent transition to the mono-allelic state occurs because it takes place after implantation into the uterus. In the cXR-tXA model bi-allelic *Xist* expression would be the consequence of reduced gene silencing under two alternative conditions. In one scenario, the absence of silencing would result in persistent bi-allelic *Xist* expression, if cXR is either not yet expressed or not yet competent to repress *Xist* (Fig. 3D+S2 left, day 1-4). The transition to the mono-allelic state would then be induced by cXR up-regulation or activation concomitantly with establishment of *Xist’s* silencing capacity (day 5-8). In a second scenario, tXA would completely resist *Xist*-mediated silencing, while cXR would be partially silenced (“dampened”, Fig. 3D+S2 right). Under these conditions, the model would predict bi-allelic *Xist* up-regulation (day 1-4), which would be resolved to a mono-allelic state upon onset of complete silencing (day 5-8). In summary, the cXR-tXA model can reproduce the different degrees of transient bi-allelic expression observed across mammals and predicts that the underlying network can resolve bi-allelic expression to the mono-allelic state that is found in somatic cells.

### Bi-allelic *Xist* up-regulation is reversible

To test the model in a more direct manner, we reasoned that by increasing the switch-ON-to-silencing ratio, we would be able to reduce the extent of transient biallelic *Xist* expression (cp. Fig. 3A). We tested this prediction experimentally in 2i-cultured mouse embryonic stem cells (mESC), which undergo random XCI upon differentiation *in vitro* with passing through a phase of bi-allelic *Xist* up-regulation^26^. The cell line used (TX1072), which was derived from a cross between two highly polymorphic mouse strains (C57BL6/J x Cast/EiJ), herein referred to as B6 and Cast, respectively, carries a doxycycline-inducible promoter upstream of *Xist* on the B6 X chromosome (Fig. 4A, top), such that *Xist* up-regulation can be accelerated by doxycyline treatment^27^. As predicted by the model (Fig. 4B), doxycycline addition one day before the onset of differentiation significantly reduced bi-allelic *Xist* up-regulation from ~20% in control cells to <5% in doxycyline treated cells (Fig. 4C). Speeding up *Xist* up-regulation can thus indeed modulate the extent of bi-allelic *Xist* expression.

The second prediction we aimed to test was that transient bi-allelic expression could be resolved to a mono-allelic state. We set up an experimental system to artificially increase bi-allelic *Xist* up-regulation. To this end, we deleted the *DXPas34* enhancer of *Tsix* from the wildtype Cast X chromosome in the TX1072 ESC line (Fig. 4D), which should result in preferential *Xist* up-regulation from that chromosome^7^. After 48h of differentiation we added doxycycline to induce *Xist* also from the other allele (B6) (Fig. 4E), thus increasing the amount of bi-allelically expressing cells from ~10% to ~30% (Fig. 4F). Since *Xist* expression from the B6 chromosome is maintained by doxycycline, the cells are predicted to down-regulate *Xist* from the Cast chromosome to resolve the bi-allelic expression state (Fig. 4G left, light green). To quantify *Xist* expression in an allele-specific way, we used the Illumina targeted expression assay to perform amplicon-sequencing of SNPs on cDNA, making use of the large number of polymorphisms present in a B6xCast cross. As predicted, *Xist* from the Cast chromosome was significantly down-regulated 48h after doxycycline treatment compared to the untreated control (Fig. 4G, right, light green). To distinguish whether *Xist* up-regulation had indeed been reversed, we performed Immuno-RNA FISH for *Xist* and H3K27me3, which is recruited to the chromosome following *Xist* RNA coating^1^. 48h after doxycycline treatment we indeed identified chromosomes that had ceased to express *Xist*, but were still enriched for H3K27me3 (as this mark is lost more slowly from the X chromatin), and named these Xa* (scheme in Fig. 4E, example image Fig. 4H). Quantification of cells that had reverted from a bi-allelic to a mono-allelic state (Xa*Xi) revealed that these were rarely observed after 4 days of differentiation without doxycycline (<5%), but constituted >10% of cells upon bi-allelic induction of *Xist* (Fig. 4I). We thus conclude that the bi-allelic *Xist* expression state can indeed be resolved by down-regulation of one *Xist* allele.

**Figure 4:**
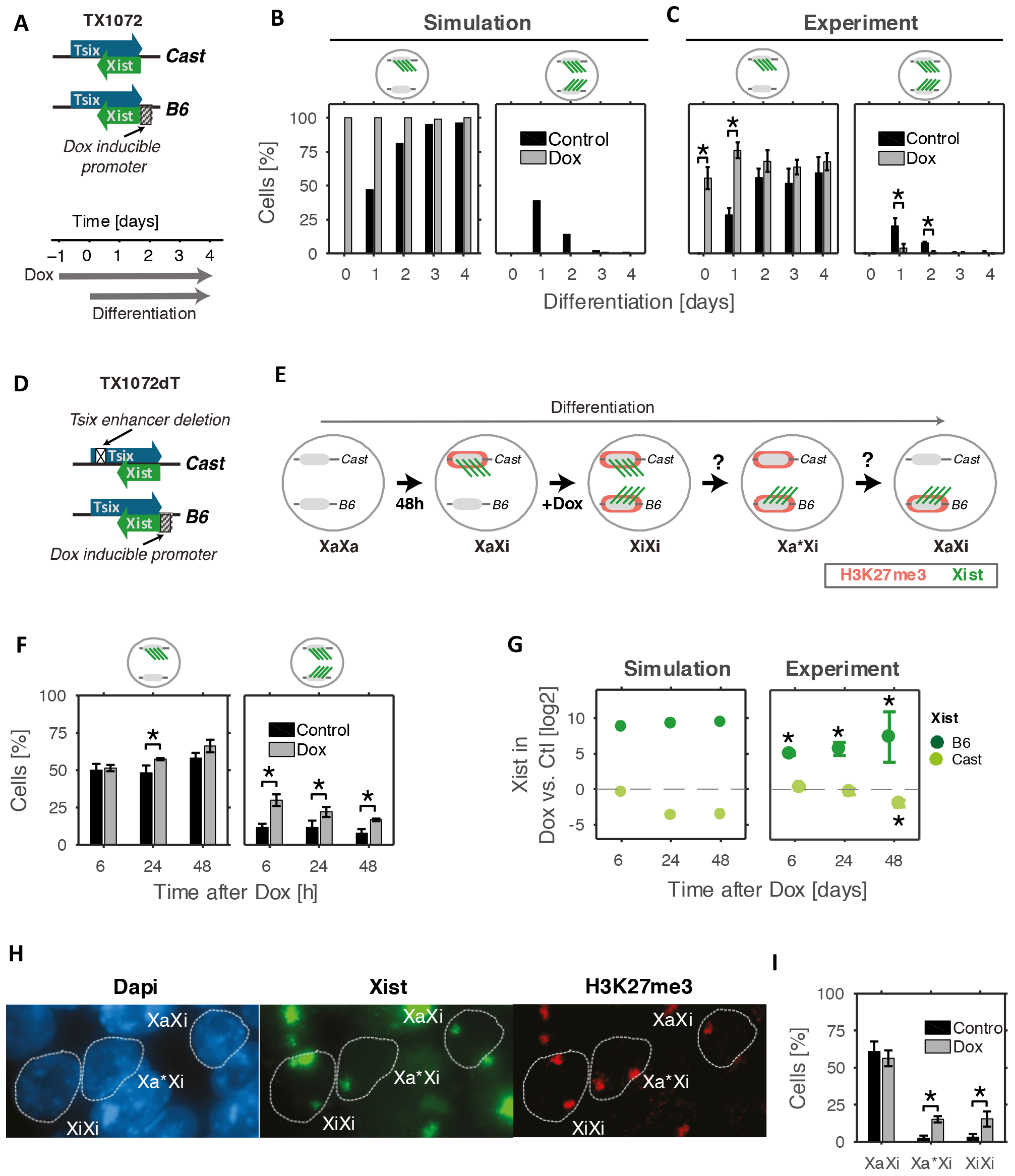
Bi-allelic *Xist* up-regulation can be resolved. **(A)** Schematic representation of the cell line used (top) and treatment performed (bottom) in (B). (**B-C)** In a simulation (B) and in an experiment (C) cells were treated with doxycycline one day before differentiation and the percentage of cells showing mono-allelic (left) and bi-allelic *Xist* up-regulation (right) is shown. **(B)** The simulation for one example parameter set is shown (the results for all tested parameter sets can be found in Fig. S3A). **(C)** *Xist* patterns were assessed by RNA-FISH (n>80). **(D-I)** Bi-allelic *Xist* up-regulation is artificially induced by treating TX1072dT cells (D) with doxycycline after 48h of differentiation. **(E)** The model predicts *Xist* down-regulation from the Cast chromosome, potentially transitioning through an Xa* state, where H3K27me3 (red) is still enriched, while *Xist* (green) has already been down-regulated. **(F)** The *Xist* expression pattern at different time points after doxycyline addition as assessed by RNA FISH (n>100). **(G)** *Xist* expression levels from the B6 and Cast alleles at different time points after doxycyline treatment as predicted by the simulation (left) and measured experimentally by Illumina targeted expression assay (right). In the simulation one example parameter set is shown, results for all other tested parameter sets can be found in supplemental Fig. S3B. **(H-I)** Immunofluorescence followed by RNA-FISH to detect *Xist* and H3K27me3 48h after doxycycline induction (n>120). **(I)** Three states were quantified (XaXi, Xa*Xi, XiXi) as shown in the example image (H). Mean and standard deviation of 3 independent experiments are shown. *p>0.05 in two-sample (C,F,I) or one-sample (G) two-sided T-test

### A mechanistic cXR-tXA model of murine *Xist* regulation

The identification of the regulator classes required for mono-allelic *Xist* up-regulation paves the way to uncovering the molecular identity of cXR and tXA. So far not tXA factor with all required characteristics has been identified (see discussion for details). As cXR candidates several *cis*-acting repressors have been proposed, such as *Linx* and *Tsix* (Table 1). *Tsix* transcription is known to be required to repress *Xist*, so that silencing of *Tsix* during X inactivation is expected to inhibit its regulatory activity on *Xist*. Mutual inhibition between *Xist* and *Tsix* seem to have the required characteristics to actually be the predicted *cis*-acting double negative feedback loop.

To test whether antisense transcription-mediated repression could generate the required bistability *in cis*, we developed a detailed mechanistic model of the *Xist/Tsix* locus, describing transcriptional initiation, RNA Pol II elongation and RNA degradation of this antisense gene pair (Fig. 5A). The model assumes three mechanisms for mutual repression of *Xist* and *Tsix*: (1) *Xist* RNA-dependent silencing of the Tsix promoter, (2) Tsix-transcription dependent repression of the *Xist* promoter and (3) transcriptional interference (TI), occurring when Pol II complexes transcribing on opposite directions meet. Since two Pol II complexes probably cannot bypass each other^28^, we assumed that one randomly chosen Pol II will be removed from the locus (for details see supplemental model description). Through a multi-step simulation process we could indeed identify parameter sets that reproduced random mono-allelic Xist up-regulation (example in Fig. 5B+C). To understand which of the inhibitory mechanisms were actually required, we developed 6 reduced models with only one or two inhibitory mechanisms. While two of the reduced models ([1,2], [1,3]) were able to maintain the XaXi state, only one of them [1,3], which retained Xist-dependent silencing and transcriptional interference (TI) could up-regulate *Xist* in a mono-allelic fashion (Fig. 5D). This reduced model was employed in all further simulations and is referred to as the “antisense model”.

**Figure 5:**
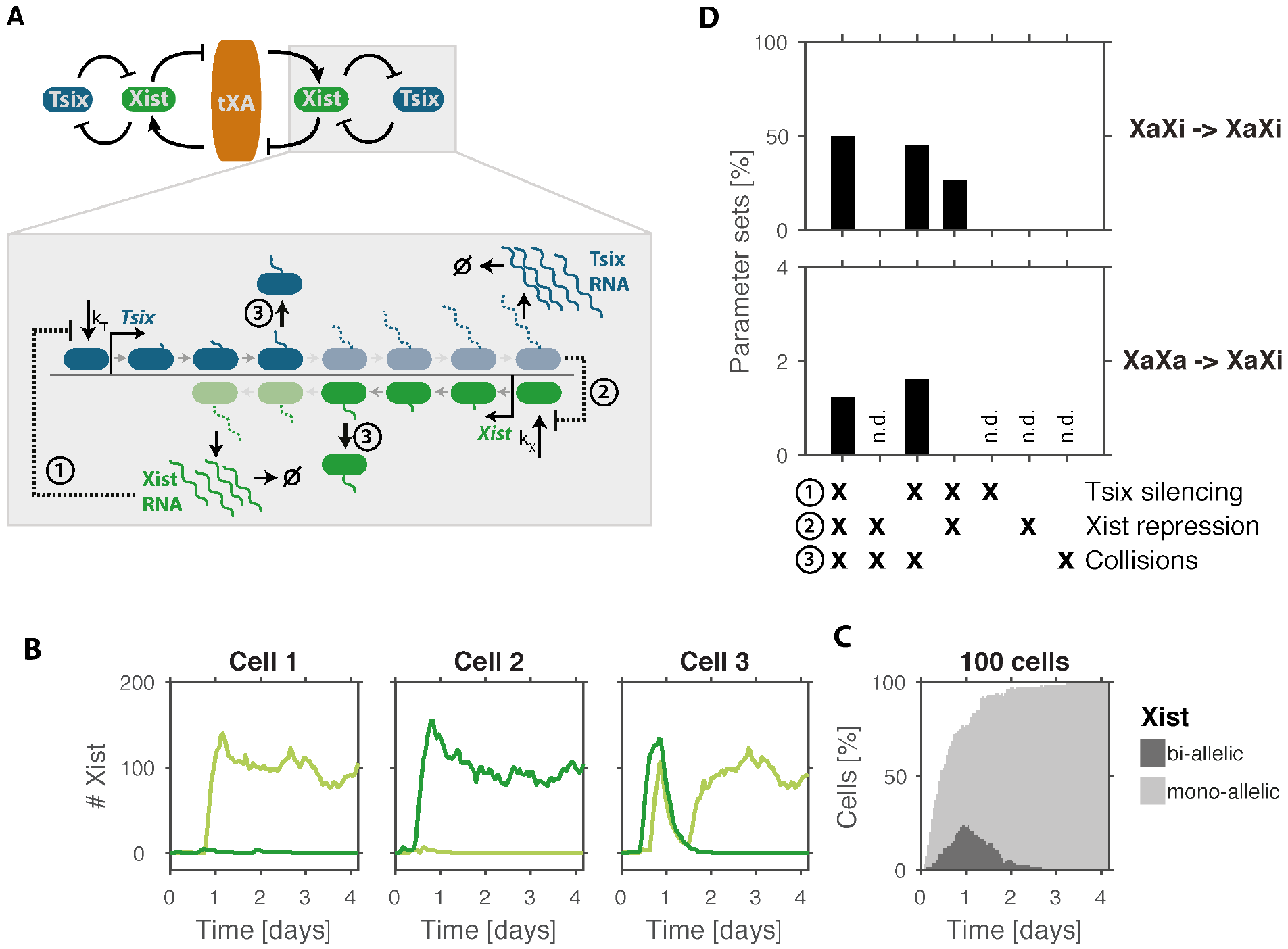
Predicted local feedback can be mediated by antisense transcription. **(A)** Schematic representation of the model were *Tsix* acts as the predicted *cis*-acting repressor: RNA Pol II complexes can bind to the *Tsix* (blue) and *Xist* (green) promoters and then move along the gene in a convergent fashion. When reaching the end of the gene they will produce an RNA molecule. Mutual repression occurs at three levels: (1) Silencing of the *Tsix* promoter by *Xist* RNA, (2) repression of the *Xist* promoter by antisense transcription and (3) random removal of one Pol II complex, if two antisense Pol II complexes occupy the same DNA element. **(B-C)** Stochastic simulation of *Xist* up-regulation for one example parameter set for the model shown in (A), showing three individual cells (B) and a population of 100 cells (C). Light and dark green in (B) represent *Xist* levels expressed from the two X chromosomes, light and dark grey in (C) represent mono- and bi-allelic *Xist* expression, as indicated. **(D)** Testing of model simplifications for the network in (A), where *Xist* and *Tsix* interact through one or two of the three repressive mechanisms as indicated. The percentage of parameter sets that can maintain the XaXi state (top) and that can initiate mono-allelic *Xist* up-regulation (bottom) in a stochastic simulation for each model are shown. Mono-allelic up-regulation was only tested for parameter sets that could maintain the XaXi state (others n.d.).

### Transcriptional interference at the *Xist/Tsix* locus

To test whether TI could indeed be observed experimentally, we assessed whether forced *Xist* transcription would interfere with *Tsix* elongation. To this end we used several ESC lines carrying the TX allele, where the endogenous *Xist* gene is controlled by a doxycycline-inducible promoter (Fig. 6A). Through doxycyline induction, *Xist* regulation can be uncoupled from the transcriptional activity of *Tsix*. Upon *Xist* induction in female TX1072 cells and an XO subclone of that line^27^ we quantified *Tsix* RNA produced by the TX allele by pyrosequencing, which allows allele-specific analysis, and by qPCR, respe ctively (primer positions in Fig. 6B). In both cell lines, *Tsix* upstream of the overlapping region (5′) was barely affected by *Xist* induction (Fig. 6C, dark blue), while *Tsix* levels downstream of *Xist* (3′) were reduced by ~50% after 8 h of doxycycline treatment (light blue). In addition spliced *Tsix* was also strongly reduced, since the splice acceptor site is close to the 3′ end (Fig. 6C, purple). These results suggest that *Xist* induction indeed interferes with *Tsix* elongation.

To further validate TI at the *Xist/Tsix* locus, we used quantitative microscopy to measure nascent transcription at the single cell level by RNA FISH with intronic oligonucleotide-based probes in a male mESC line carrying the TX allele (TXY)^29^. For *Tsix* we designed two different probes to detect transcription upstream of *Xist* (5′) and within the overlapping region (3′) (Fig. 6B). Transcription of *Xist* and *Tsix* was mutually exclusive in nearly all cells after one day of doxycyline treatment (Fig. 6D). To be able to observe transcriptional interference independent of *Xist* RNA mediated silencing, we used the silencing deficient TXYΔA line carrying a deletion of the A-repeat of *Xist^29^*. Strikingly, almost complete *Tsix/Xist* exclusion in the overlapping region was observed also in this case (Fig. 6E). We next compared the signal intensity of the two *Tsix* probes at *Xist* transcribing (*Xist*+) and not transcribing (*Xist*−) alleles. In the TXY line both *Tsix* signals of the 3′ and 5′ probes were strongly reduced on the *Xist+* alleles compared to *Xist*− alleles likely due to *Xist*-RNA mediated silencing of *Tsix*, while in TXYΔA cells the *Tsix* 5′ region was barely affected compared to the 3′ position (Fig. 6F). In summary, transcriptional interference perturbs transcriptional elongation at the *Xist*/*Tsix* antisense locus, thus validating a central assumption of the antisense model.

### The antisense model can reproduce *Xist* and *Tsix* mutant phenotypes

To further validate the antisense model, we tested whether it could reproduce known *Xist* and *Tsix* mutant phenotypes. For 100 parameter sets that could reproduce mono-allelic Xist up-regulation (cp. Fig. 5) four genotypes were simulated: wildtype, heterozygous and homozygous *Tsix* mutant and heterozygous *Xist* mutant cells (Fig. 7A-D).

**Figure 6:**
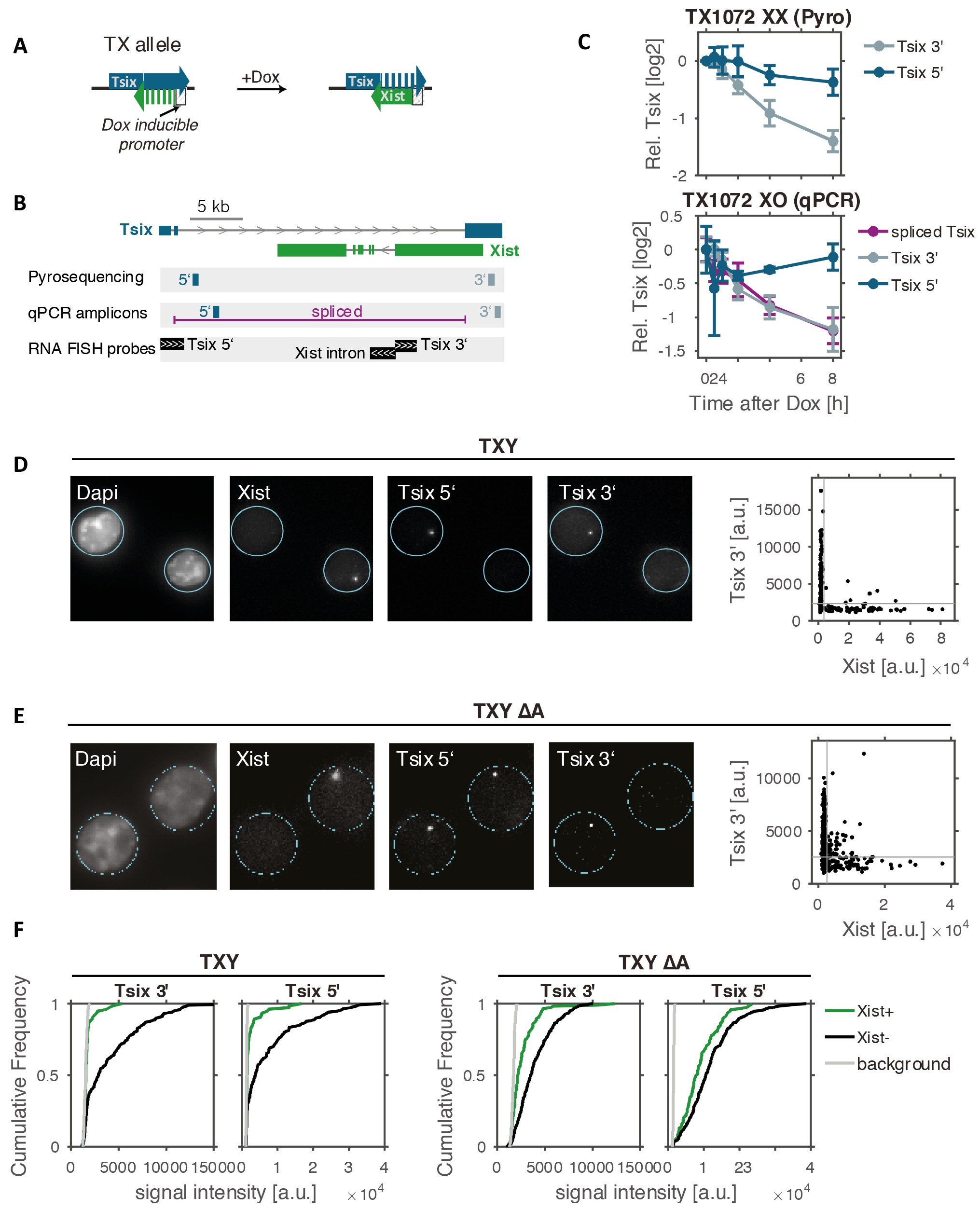
Transcriptional interferences at the *Xist/Tsix* locus. **(A)** The TX allele carries a doxycycline-inducible promoter driving the endogenous *Xist* gene and was used to investigate whether *Xist* transcription would interfere with Tsix elongation in (C-F). **(B)** Position of primers and probes used in (C-F). **(C)** TX1072 XX (top) and TX 1072 XO ESCs (bottom) were treated with doxycycline for 8 h and *Tsix* transcription from the TX allele was assessed by pyrosequencing (XX) or qPCR (XO) at different positions within the *Tsix* gene. Mean and standard deviation of three independent experiments are shown. **(D-E)** TXY (D) and TXY ΔA (E) ESCs were treated with doxycycline for 24 hours and nascent transcription of *Xist* and *Tsix* (5′ and 3′) was assessed by RNA FISH (probe positions in B). Example images and quantification (right) of 245 (D) and 422 cells (E) is shown, where each dot represents the measured signal intensities of a single allele/cell. Grey lines indicate the detection threshold estimated from negative control regions. **(F)** Cumulative distribution of *Tsix* signal intensity at *Xist+* (green) and *Xist*− alleles (black) in TXY (left) and TXYΔA (right). For TXYΔA one out of 3 experiments is shown.

In our simulations, each chromosome in wildtype cells is inactivated with an equal probability such that 50% of cells will express *Xist* from one or the other X chromosome (Fig. 7A, bottom). In agreement with experimental observations^7^, a heterozygous *Tsix* mutation results in non-random X inactivation of the mutant X chromosome (Fig. 7B, bottom). A heterozygous *Xist* deletion by contrast results in complete skewing towards the wildtype allele, but nevertheless most cells up-regulated *Xist* after 4 days of differentiation (Fig. 7D, bottom). For homozygous *Tsix* mutants, ‘chaotic’ X inactivation has been described with a mixture of cells inactivating one or two X chromosomes^30^. In our simulations such a mutation would result in *Xist* oscillations, where *Xist* is turned on bi-allelically, which will result in complete tXA silencing and *Xist* down-regulation, followed by another round of bi-allelic up-regulation (Fig. 7C, top). In agreement with the experimental phenotype, these simulations show a strongly increased fraction of bi-allelically expressing cells (Fig. 7C, bottom).

We also compared the kinetics of *Xist* up-regulation in the mutants, since in a previous study a heterozygous *Tsix* mutation was shown to accelerate XCI, while an *Xist* mutation was found to slow down X inactivation^16^. To test this, we calculated the half time of *Xist* up-regulation (T_1/2_) where 50% of cells would have turned on *Xist* (example in Fig. 7E) and compared this value between mutant and wildtype cells. Indeed for all parameter sets tested, a *Tsix* mutation would reduce the half time of *Xist* up-regulation while it would be increased in *Xist* mutant cells (Fig. 7F). Taken together, these results strongly support antisense-mediated repression of *Tsix* as a promising candidate for the predicted cXR factor in mice.

## Discussion

Through screening 36 alternative network architectures, we have identified a core network that can recapitulate random mono-allelic and female-specific *Xist* up-regulation. This network, consisting of a *trans*-acting activator and a *cis*-acting repressor, resembles an “extended toggle-switch”, which is thought to govern many cell fate decisions through generating mutual exclusive expression of antagonizing lineage specifying transcription factors^31^. While two transcription factors on factors such as PU.1 and Gata1 (driving myeloid and lymphoid differentiation, respectively) mutually repress each other in a classical toggle switch^31^, in our network the two copies of *Xist* inhibit each other though silencing of the *trans*-activating tXA factor. However, this inhibition cannot be directional, since both *Xist* loci will be affected by reduced levels of any *trans*-acting regulator. To nevertheless allow the establishment of two alternative states in such a symmetric network, we have shown here that a local positive feedback, mediated by the *cis*-repressor cXR, is required to memorize the initial choice of the inactive X, at least until the two states could be locked in by epigenetic mechanisms such as DNA methylation^1^. During cell fate decisions, transcription factors often promote their own expression though similar positive feedback regulation. Cells thus seem to employ strikingly similar regulatory principles to ensure mono-allelic *Xist* up-regulation, as in other unrelated molecular decision-making processes.

**Figure 7:**
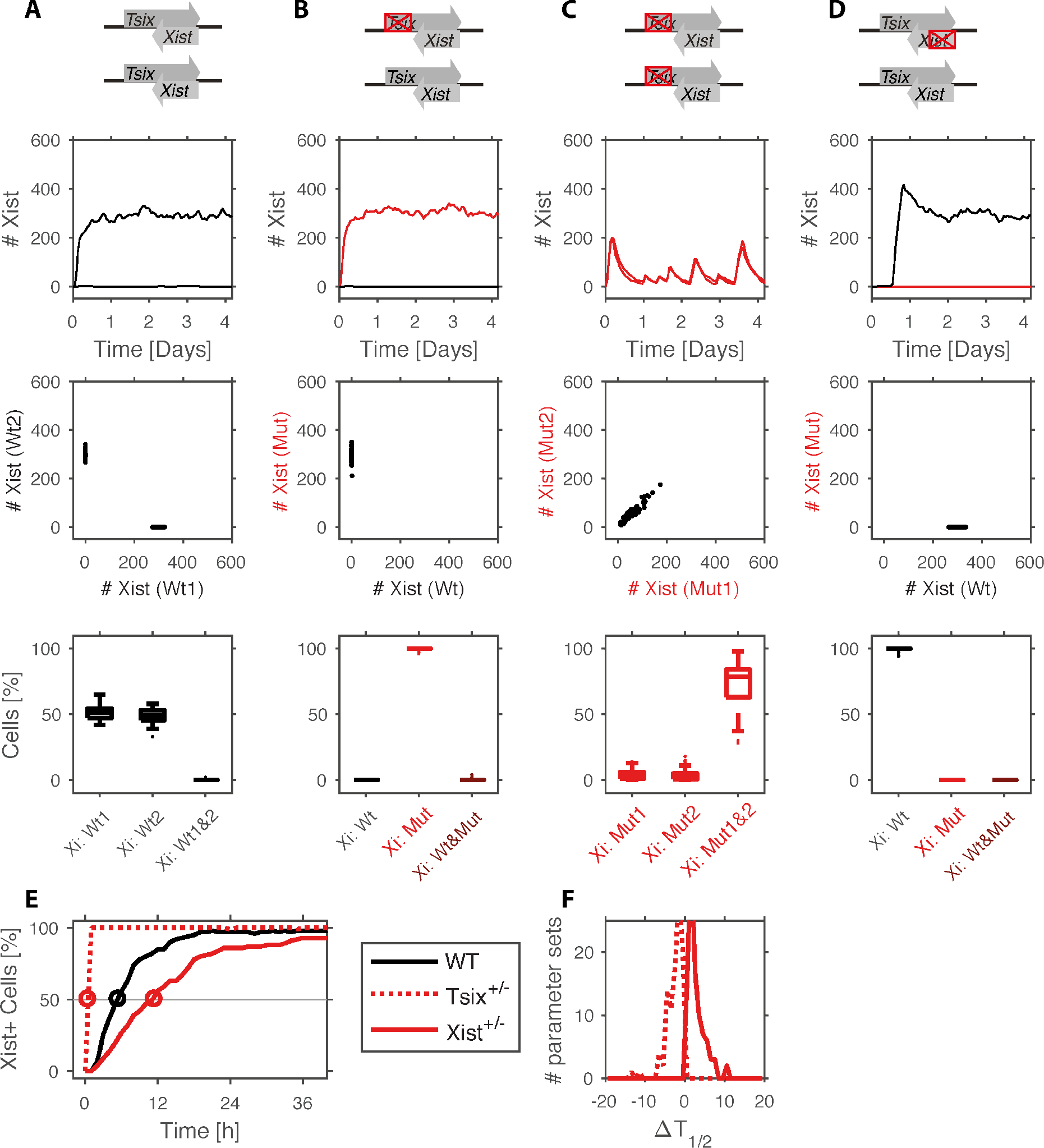
*Xist/Tsix* mutant simulations. **(A-D)** Simulations of *Xist/Tsix* mutant cell lines (top). Representative simulation of *Xist* levels produced by the wildtype (black) and mutant (red) chromosomes of a single cell (upper middle) and by 100 cells (lower middle). Boxplots (bottom) show the percentage of cells expressing *Xist* mono-allelically from wildtype or mutant X or bi-allelically for all 100 simulated parameter sets. On each box, the central mark indicates the median, and the bottom and top edges of the box indicate the 25th and 75th percentiles, respectively. The whiskers extend to the most extreme data points not considered outliers, and the outliers are plotted individually. **(E-F)** *Xist* up-regulation is accelerated in *Tsix*^+/−^ and delayed in *Xist*^+/−^ cells. Representative simulation (E) and the distribution of the change of half time (ΔT_1/2_) in the mutant genotypes (F).

The generic network we have identified will serve as a framework to uncover the molecular identity of the required regulators. Our approach is thus highly complementary to previous studies that have identified and characterized individual regulators of *Xist*, usually one at a time^13–15,32^. While the known X-linked regulators can be grouped according to the classification we have developed (Table 1), autosomal factors are not explicitly accounted for in our modeling framework. They might modulate reaction rates in a differentiation dependent manner (e.g. pluripotency factors^33–35^) or mediate the effects of X-linked regulators (e.g. Rex1 as a target of Rnf12^19^). X-linked regulators outside the identified core network might add additional robustness to the system (e.g. *Jpx*) and/or mediate interactions within the core network (e.g. Ftx could be targeted by a tXA factor).

So far, two *trans*-acting activators of *Xist* have been proposed (Table 1), the E3 ubiquitin ligase Rlim/Rnf12, which targets the *Xist* repressor Rex1/Zfp42 for degradation, and the lncRNA *Jpx*^13,19,36^. While *Jpx* escapes XCI, Rnf12 is rapidly silenced by *Xist* and has thus been suggested to form the *trans*-acting negative feedback loop that we have also identified through our network screening approach^15^. Rnf12 could thus be the tXA factor that our analysis predicts. However, while overexpression of Rnf12 can induce *Xist* ectopically in male cells, its deletion in females does not necessarily prevent *Xist* up-regulation^15,18,37,38^, suggesting that additional tXA factors remain to be identified.

The cXR factor is likely a lncRNA locus, since this class of regulators frequently act *in cis*^39^. In mice, two such loci, *Tsix* and *Linx/Ppnx* have been described^12,32^. Since transcription seems to be dispensable for the function of *Linx* (R. Galupa, E. Heard, in preparation), it is probably insensitive to *Xist*-mediated silencing and would therefore not be expected to form a feedback loop. *Tsix*, by contrast, must be transcribed to exert its repressive function by depositing repressive histone marks at the *Xist* promoter during antisense transcription^40,41^. By developing a mechanistic model of the *Xist/Tsix* locus, we show that mutual repression mediated by antisense transcription can indeed generate a local switch. Through transcriptional interference, which we confirmed to occur experimentally, antisense transcription can generate a very precise threshold (data not shown) and thereby ensure reliable mono-allelic *Xist* up-regulation. While the function of *Tsix* in mice is well documented, its conservation in other mammals, such as humans has not been shown^2^. Human *TSIX* has so far neither been detected in embryonic stem cells nor in embryos. Its transcription has been reported in embryoid body derived cells of rather undefined origin, where is was found to be truncated compared to mouse *Tsix* and appeared to be co-expressed with *XIST* from the same allele^8^. However, since the establishment of random XCI has not yet been observed experimentally neither *in vivo* nor *in vitro*, it might still be accompanied by *TSIX* transcription^42^. Even with the reduced overlap between *XIST* and *TSIX* reported for the human locus, the TI-based switch assumed in our antisense model could in principle still ensure mono-allelic Xist expression (data not shown). The functional conservation of *TSIX* in humans therefore still remains an open question. Interestingly, another lncRNA, *XACT*, appears to antagonize *XIST in cis* specifically in humans and could thus function as a cXR factor in that species^9^. Alternatively, the *cis*-acting positive feedback loop could also be implemented at the level of chromatin readers and writers as shown for the *flc* locus in Arabidopsis^43,44^.

While the precise implementation of the predicted positive feedback might vary between different mammals, the basic network structure we have identified can recapitulate all expression patterns that have been observed in mice, humans and rabbits. Depending on the relative time scales of *Xist* up-regulation and gene silencing, the same network can produce different degrees of transient bi-allelic expression and thereby recapitulate both low and high levels of bi-allelic *Xist* up-regulation as observed in mice and rabbit embryos, respectively^3–5^. Through ectopic induction of bi-allelic *Xist* expression we show that this state is reversible during early differentiation. Our model predicted that some level of bi-allelic up-regulation should also occur in mice, if initiation of XCI is to be completed within a physiologic time frame. When analyzing mouse embryos right at the onset of random XCI at E5.0, we indeed observed ~20% bi-allelic *Xist* expression, which is in agreement with another recent study^25^.

The situation in human pre-implantation embryos seems to be special in the sense that *Xist’s* silencing ability is reduced or even absent^3,6^, possibly because factors that mediate silencing are not expressed at these stages of development. The core network we have identified would predict extended bi-allelic *Xist* expression (as observed in human embryos^3^) to arise from reduced gene silencing if either (1) cXR was not yet expressed or active at this stage or (2) cXR was dampened, while tXA was insensitive to *Xist*. Establishment of *XIST’s* silencing capacity (together with cXR up-regulation/activation in scenario 1) would be predicted to induce a transition to the mono-allelic state. In particular scenario 1 is intriguing since antisense transcription, which appears to function as cXR in mice, is not observed during human pre-implantation development, but could potentially be up-regulated when the transition to the mono-allelic state occurs. If cXR was already expressed, but not yet able to repress *XIST*, it would be co-expressed with *XIST* on both chromosomes first, but restricted to the active X after XCI, a pattern highly reminiscent of *XACT* in pre-implantation embryos and post-XCI human ESCs, respectively. Although so far the onset of random XCI could not be recapitulated with human ESCs^42^, further refinement of the culture conditions will hopefully allow us to test whether indeed mono-allelic *XIST* expression is established once silencing sets in and whether this might be accompanied by antisense transcription. Taken together, our study reveals that the regulatory principles employed by different mammalian species might be less diverse than previously thought and that the different routes to the mono-allelic state could be attributed to quantitative differences in reaction rates rather than qualitative differences in the network architecture.

## Methods

### ODE simulations

ODE models were formulated by assuming a hill type regulation of production rates and first-order degradation rates, in a way that the levels of all variables were scaled between 0 and 1. Interaction between two regulators was assumed to occur synergistically. Simulations were performed in Matlab using the ode15s integrator. Details are given in the supplemental model description.

### Stochastic simulations of the cXR-tXA model

Reactions describing production and degradation reactions were formulated directly from the ODE model, with adding scaling factors that would determine the maximal number of molecules that can be produced for a given variable. Moreover, each *Xist* molecule had to transition through a certain number of silencing intermediates (with rate 1h^−1^) before reaching a silencing-competent state. The silencing delay for cXR (sil_cXR_) and tXA (sil_cXR_) was then given by the number of required silencing intermediates and is equal to the mean silencing delay. The simulations were performed using the Gillespie algorithm^45^, implemented in Julia and executed on a computing cluster.

### Stochastic simulations of the antisense model

To simulate antisense transcription, RNA Pol II molecules were assumed to bind to the *Tsix/Xist* promoters in a stochastic fashion and then moved along the *Xist/Tsix* locus deterministically, which had been divided into 100nt long segments. For elongation and degradation rates experimental estimates from the literature were used^36,46^. When *Xist* RNA exceeded a threshold of 10 molecules, the *Tsix* promoter could switch to the OFF state. Transcription of a *Tsix* polymerase through the *Xist* promoter, induced a switch to the OFF state, that could revert back to the ON state with a constant rate. When two RNA Pol II molecules occupied the same DNA segment, one randomly chosen polymerase was removed from the gene. tXA produced per allele was scaled between 0 and 1 and was set to 0 with a certain delay after *Xist* had exceeded a threshold of 10 molecules. Simulations were conducted in MATLAB. The model was written in C++ and compiled into a MEX file that was called from the main MATLAB function. For parameter scanning a compiled Matlab script was executed in parallel on a computing cluster.

### Cell lines

The female TX1072 cell line and its subclone TX1072 XO (clone A11) are F1 hybrid ESCs (CastxB6), which carry a doxycycline responsive promoter in front of the *Xist* gene on the B6 chromosome and an rtTA insertion in the Rosa26 locus (described in ^27^) The TX1072dT line (clone 1C6) was generated by introducing a deletion of the Dxpas34 repeat in TX1072 cells on the Cast chromosome by co-transfecting Cas9 expression vectors p330 expressing sgRNAs GTACATAATGACCCGATCTC and GAACTCACTATATCGCCAAAG ^47^. Clones with the deletion were identified by PCR (ES585:AGGCACACCACCCCAGTGGA, ES609:TCCAAACATGGCGGCAGAAGC) and the deleted allele was identified by Sanger sequencing of the PCR product using primer ES609 based on two SNPs at positions 100,645,601 (Cast: C) and 100,641,221 (Cast: G) (mm9). Male-inducible wild-type and ΔA *Xist* lines were a gift from A. Wutz (called *Xist*-tetOP and *Xist*-ΔSX-tetOP, respectively, in ^29^)

### ES cell culture and differentiation

TX1072, TX1072 XO and TX1072dT(clone 1C6) cells were grown on gelatin-coated flasks in serum-containing ES cell medium (DMEM (Sigma), 15% FBS (Gibco), 0.1mM β-mercaptoethanol, 1000 U/ml leukemia inhibitory factor (LIF, Chemicon)), supplemented with 2i (3 μM Gsk3 inhibitor CT-99021, 1 μM MEK inhibitor PD0325901) for TX1072 and TX1072 XO. Differentiation was induced by 2i/LIF withdrawal in DMEM supplemented with 10% FBS and 0.1mM β-mercaptoethanol at a density of 4*10^4^cells/cm^2^ in Fibronectin (10 μg/ml) coated tissue culture plates. For ectopic *Xist* induction the medium was supplemented with 1 μg/ml Doxycycline. To induce *Xist* in undifferentiated cells, they were plated at a density of 1*10^5^cells/cm^2^ two days before the experiment and then treated with 1 μg/ml Doxycycline. Male-inducible wild-type and ΔA *Xist* lines were grown on mitomycin-C-inactivated mouse embryonic fibroblasts in ES cell media containing 15% FBS (Gibco), 0.1mM β-mercaptoethanol (Sigma), 1,000 U/ml LIF (Chemicon) and treated for 24 h with 2 μg/ml doxycycline.

### Mice

All animal experiments were performed under the ethical guidelines for the Care and Use of Laboratory Animals (French ethical committee of animal experimentation: Institut Curie #118 and agreement C75-05-17 for the animal facility; Kyoto University). Embryos were obtained by natural mating between B6D2F1 (derived from C57BL/6J and DBA2 crosses) female and males. Noon of the day when vaginal plugs were detected was set as E0.5.

### Conventional RNA FISH on ESCs

FISH on cells from tissue culture was performed as described previously ^48^. Briefly, mESCs were dissociated using Accutase (Invitrogen) and adsorbed onto Poly-L-Lysine (Sigma) coated coverslips #1.5 (1mm) for 5 min. Cells were fixed with 3% paraformaldehyde in PBS for 10 min at room temperature and permeabilized for 5 min on ice in PBS containing 0.5%Triton X-100 and 2mM Vanadyl-ribonucleoside complex (New England Biolabs). Coverslips were preserved in 70% EtOH at −20°C. Prior to FISH, samples were dehydrated through an ethanol series (80%, 95%,100% twice) and air-dried quickly. For detecting Huwe1, a BAC spanning the respective genomic region (RP24-157H12) was labeled by nick translation (Abbot) using dUTP-Atto550 (Jena Bioscience). Per coverslip, 60ng probe was ethanol precipitated with Cot1 repeats, resuspended in formamide, denatured (10min 75°C) and competed for 1h at 37°C. *Xist* was detected with a custom designed strand-specific probe that covers all exons with ~75bp long oligo nucleotides end-labeled with the Alexa488 fluorophore (Roche). Both probes were co-hybridized in FISH hybrization buffer (50% Formamide, 20% Dextran sulfate, 2x SSC, 1μg/μl BSA, 10mM Vanadyl-ribonucleoside) over night. Washes were carried out at 42°C three times 7min in 50% formamide in 2X SSC at pH=7.2 and three times 5min in 2X SSC. 0.2mg/mL DAPI was used for counterstaining and mounting medium consisted in 90% glycerol, 0.1X PBS, 0.1% p-phenylenediamine at pH9 (Sigma). Images were acquired using a wide-field DeltaVision Core microscope (Applied Precision) or a widefield Z1 Observer (Zeiss) using a 100x objective.

### Immunofluorescence combined with RNA FISH

For immunofluorescence staining cells were differentiated on Fibronectin coated cover slips (18mm Marienfeld) at a density of 2*10^4^cells/cm^2^. Cells were fixed and permeabilized as described above and incubated with the H3K27me3 antibody (Active Motif #39155, 0.4ug/ml) in PBS for 1h at room temperature, then washed 3 times for 10 minutes with PBS, followed by a 1h incubation with an Alexa-555 labelled Goat anti-rabbit antibody (Invitrogen A-21428, 0.8 ug/ml). After 3 washes, the cells were fixed again with 3% paraformaldehyde in PBS for 10 min at room temperature, followed by three short washes with PBS and two washes with SSC. Hybridization was then performed as described in above.

### Quantitative RNA FISH

Quantitative RNA FISH on *Xist* and *Tsix* was performed using Stellaris FISH probes (Biosearch Technologies). Cells were adsorbed and fixed as described above. Cells were prehybridized in wash buffer (2x SSC, 10% formamide) twice for 5 min, then hybridized with a solution that contained 125 nM of each FISH probe, 2X SSC, 10% formamide, 10% dextran sulfate overnight at 37 °C. Cells were washed twice with wash buffer for 30 min before counterstaining DNA with 0.2mg/ml DAPI in 1x PBS, and mounted on slides using the mounting medium described above. Z-stacks were acquired using a wide-field DeltaVision Core microscope microscope equipped with a 100x objective (voxel size 129×129×200 nm). Quantification of nascent RNA signals was performed as in^49^. Briefly, the fluorescence background of each z plane was generated by morphologically opening the image with a circular structuring element with a diameter of 5 pixels (645 nm), and subtracted from the original image. A region of interest (ROI) of constant volume (30×30×6 pixels = 1.3×1.3×1.2μm) was selected around each transcription site. To reduce residual high-frequency fluorescence background, the average pixel intensity was measured in a 3-voxel thick frame adjacent to the border of the ROI, and further subtracted. The integrated intensity of the fluorescent signal was then measured within the whole ROI. Integrated intensities of approximately 200 random nuclear background ROIs were used to define a threshold to classify transcribed versus non transcribed loci.

### RNA FISH of epiblast cells from E5.0 embryos

For E5.0 mouse embryos, the embryos were dissected out from decidua and the Reichert’s membrane was removed in a 6cm Petri dish containing PBS using sharpened forceps. Extra embryonic ectoderm was separated by a fine glass needle. The epiblast/visceral endoderm were incubated in 0.25% Pancreatin (Sigma) / 0.5% Trypsin / Polyvinylpyrrolidone (PVP40; Sigma) at 4°C for 10min and transferred to a 3.5cm petri dish containing a large volume of 1%BSA/PBS. Epiblast and visceral endoderm were separated by pipetting with a mouth pipette whose internal diameter is slightly smaller than that of epiblast. RNA FISH were carried out as described previously 5, using a non strand-specific probe detecting *Xist* and *Tsix*. Embryos with an *Xist* cloud were identified as female. Images were acquired using a 200M Axiovert fluorescence microscope (Zeiss) equipped with an ApoTome was used to generate 3D optical sections. Sequential z-axis images were collected in 0.3 *μ* m steps. Images were analyzed using ImageJ software (Fiji, NIH).

### RNA extraction, reverse transcription, qPCR

For pyrosequencing and qPCR, cells were lysed by direct addition of 1 ml Trizol (Invitrogen). Then 200μl of Chloroform was added and after 15 min centrifugation (12000xg, 4°C) the aqueous phase was mixed with 700 μl 70% ethanol and applied to a Silica column (Qiagen RNAeasy Mini kit). RNA was then purified according to the manufacturer’s recommendations, including on-column DNAse digestion. For quantitative PCR (qPCR), 1ug RNA was reverse transcribed using Superscript III Reverse Transcriptase (Invitrogen). Expression levels were quantified using 2× SybRGreen Master Mix (Applied Biosystems) and a ViiA7 system (Applied biosystems) with ~8ng cDNA and the primers given in Table S1. Expression levels were normalized to Rrm2 and Rplp0.

### Targeted RNA Expression Assay

For allele-specific quantification of *Xist* expression the TruSeq Targeted RNA Expression assay (Illumina) was used according to the manufacturer’s recommendations. RNA was extracted using the Direct-zol RNA MiniPrep kit (Zymo Research) and DNase digest was performed using Turbo DNA free kit (Ambion). Four amplicons containing SNPs within *Xist* were measured and normalized to amplicons in four autosomal reference genes (Rrm2, Rplp0, Fbxo28, Exoc1). Details on the amplicons are given in supplementary table S1. Read counts of each *Xist* amplicon mapping to the B6 and Cast alleles, respectively, were normalized to the geometric mean of the four reference genes. The fold change of the doxycycline treated sample relative to the corresponding control sample was then calculated for each *Xist* amplicon. Using a one-sample t-Test it was tested whether the mean log2 fold-change of the four amplicons was significantly different from 0 (p<0.05).

### Pyrosequencing

For allele-specific expression analysis of *Tsix*, pyrosequencing technology was used. Two different amplicons within *Tsix*, each containing a SNP were PCR-amplified from cDNA with biotinylated primers and sequenced using the Pyromark Q24 system (Qiagen). Primer sequences are given in supplemental Table S1. The assay provides the fraction of *Tsix* transcript arising from the B6 chromosome at time t (Ft). To calculate the expression from the B6 chromosome at time t relative to the uninduced state at 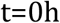 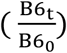 the data was transformed as follows. Assuming that expression from the Castaneus chromosome (Cast) is constant over time, 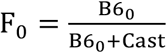 and 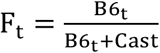 can be transformed into 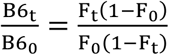

## Data Availability

The raw data produced in this study (Fig. 3b+c, Fig. 4, Fig. 5) is available upon request. The cell lines TX1072dT generated for this study is available upon request.

## Code Availability

All code used in this study is available upon request.

## Author contributions

Conceptualization, E.G.S., E.H. and I.O.; Software, V.M. and E.G.S.; Investigation, V.M., I.O., I.D., L.G., E.G.S.; Writing, E.G.S. and V.M. with input from E.H. and L.G; Supervision, E.G.S., E.H., M.S.; Funding Acquisition, E.G.S., E.H., L.G., M.S.

## Acknowledgements

We thank A. Wutz for the TXY(*Xist*-tetOP) and TXYΔA (*Xist*-ΔSX-tetOP) mESC lines. We thank the staff of the MPIMG and PICTIBiSA@BDD imaging facilities for technical assistance, the MPIMG sequencing core facility of sequencing services and the MPIMG IT for support in using the computing cluster. This work was funded by an HFSP long-term fellowship (LT000597/2010-L) to E.G.S., a grant-in-aid from MEXT and JST-ERATO to I.O. and M.S., JSPS KAKENHI Grant Number JP25291076 to I.O., the Max-Planck Research Group Leader program and and the German Ministry of Science and Education (BMBF) through the grant E:bio Module III - Xnet. V.M. is supported by the DFG (GRK1772: Computational Systems Biology).

## Supplemental Figures

**Figure S1 (linked to Fig. 2).**
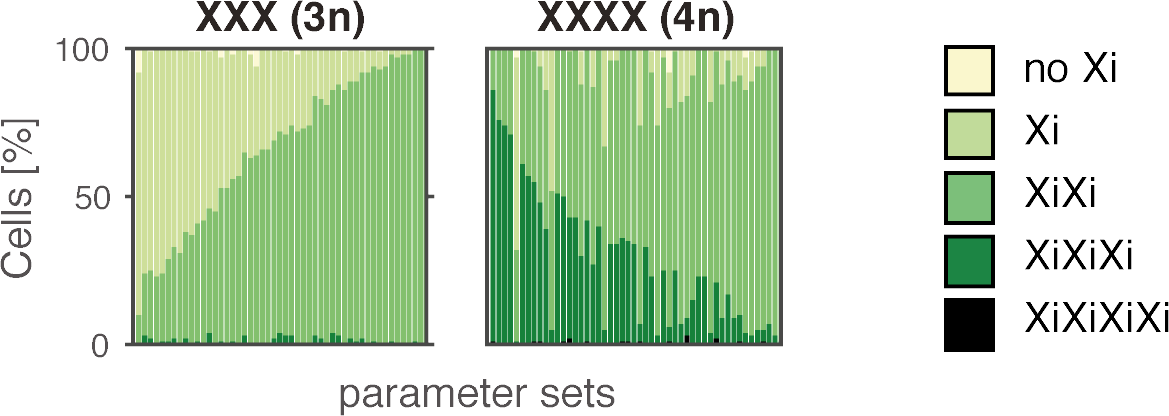
Simulations of triploid (left) and tetraploid cells (right) assuming that tXA concentrations are diluted 1.5- and 2-fold in tri- and tetraploid cells, respectively, due to an increase in nuclear volume. Stacked bar graphs show the classification of *Xist* patterns in simulations with 50 parameter sets that can reproduce mono-allelic *Xist* up-regulation in diploid cells.

**Figure S2 (linked to Fig. 3).**
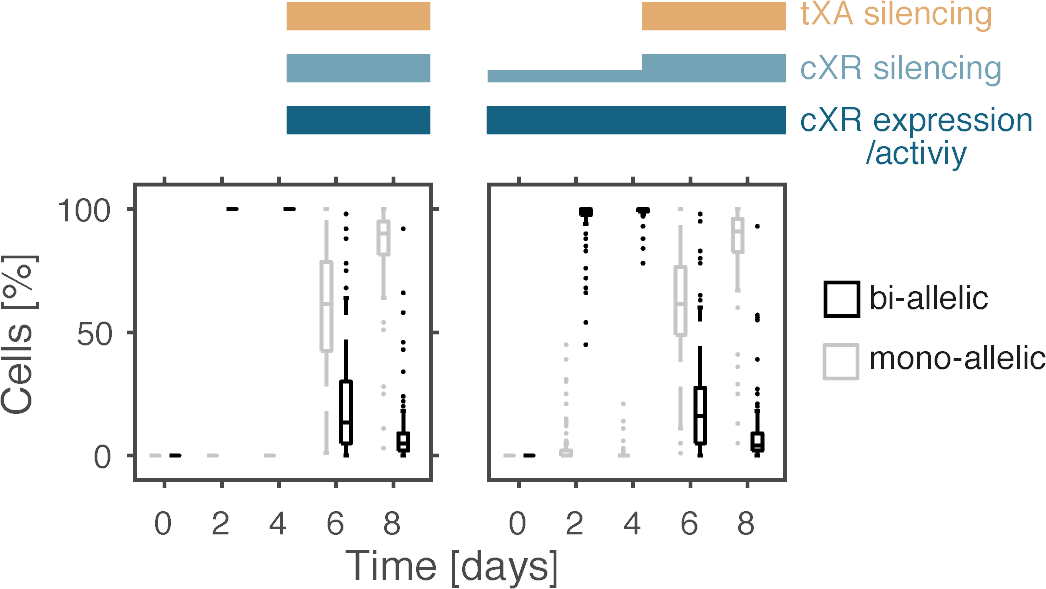
Simulation of bi-allelic expression upon reduced *Xist*-mediated silencing as observed in human embryos, assuming that in the first 4 days of the simulation either silencing and cXR expression is absent (left) or that cXR is silenced partially (dampening), while tXA is unaffected by *Xist* (right). Boxplots show the percentage of mono- and bi-allelically expressing cells for 100 randomly chosen parameter sets that could reproduce mono-allelic *Xist* up-regulation. On each box, the central mark indicates the median, and the bottom and top edges of the box indicate the 25th and 75th percentiles, respectively. The whiskers extend to the most extreme data points not considered outliers, and the outliers are plotted individually.

**Figure S3 (linked to Fig. 4).**
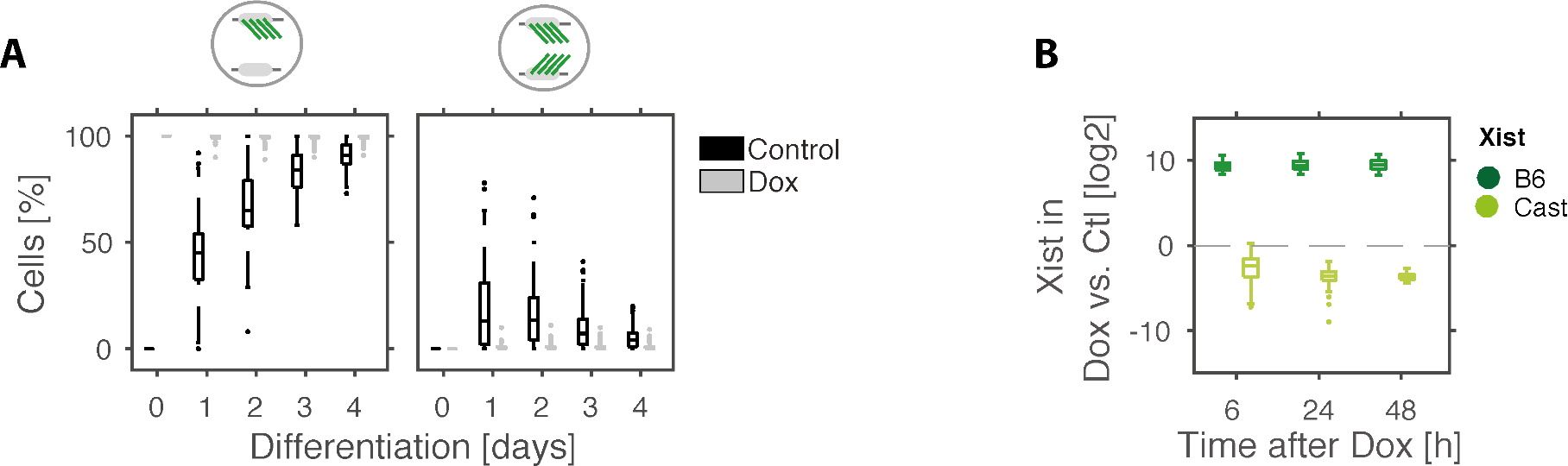
**(A)** Simulation of Doxycyline treatment one day before the onset of differentiation. Boxplots show the frequency of mono-allelic (left) and bi-allelic *Xist* expression (right) in dox-treated (grey) and control cells (black) for 100 randomly chosen parameter sets that could reproduce mono-allelic *Xist* up-regulation. **(B)** Boxplots show the simulation results for artificial bi-allelic *Xist* induction as described in Fig. 4E in the main text, using the same parameters sets as in (A). On each box, the central mark indicates the median, and the bottom and top edges of the box indicate the 25th and 75th percentiles, respectively. The whiskers extend to the most extreme data points not considered outliers, and the outliers are plotted individually.

## Supplemental Model Description

### 1 Model Comparison

#### 1.1 Single regulator models

To investigate the requirements for mono-allelic and female-specific Xist expression, all X-linked Xist regulators were grouped into 8 different classes according to whether they activate (XA) or repress (XR) *Xist*, whether they act *in cis* (c) or *in trans* (t) and whether they are silenced by *Xist* or escape XCI. To simulate regulation of *Xist* by each regulator class, 8 different ordinary differential equation (ODE) models were constructed. as follows.

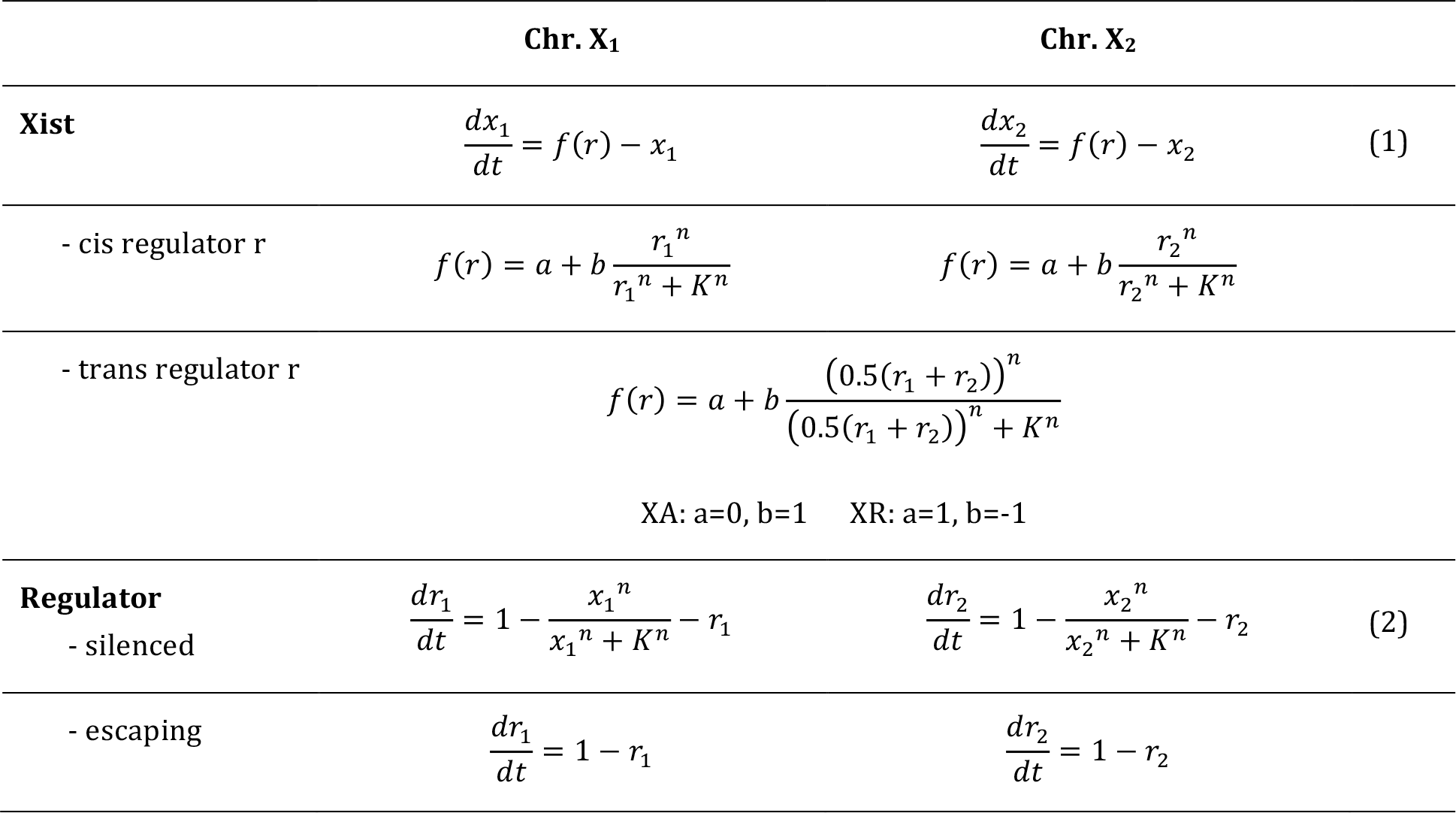

Activation or repression of *Xist* by a regulator and silencing of the regulator by *Xist* are described as hill functions with two parameters: a hill coefficient n that describes the cooperativity or threshold behavior of the interaction and a threshold K, which is the regulator level where activation or repression is half-maximal. Degradation of *Xist* and its regulators is assumed to occur with first-order kinetics with a degradation rate of 1h^−1^. Since the production rates can maximally be 1 and degradation rates are set to 1, the levels of *Xist* and its regulators can vary between 0 and 1. Since each interaction is described by two parameters, a hill coefficient n and a threshold K, each model has either 2 (escaping regulator) or 4 parameters (silenced regulator).

#### 1.2 Simulating mono-allelic expression

To assess which network can maintain stable mono-allelic expression (simulation 1), cells with one active X (Xa), where *Xist* expression is low and with one inactive X (Xi), where *Xist* expression is high were simulated with the following initiation conditions.

| Xist (Xi) | Xist (Xa) | Regulator (Xi) | Regulator (Xa) |
| --- | --- | --- | --- |
| 1 | 0.01 | 0.01 (silenced)<br>1 (escaping) | 1 |

Each network was simulated with at least 1,000 parameter sets, where parameter values were randomly drawn from a uniform distribution between 1 and 5 for hill coefficients (n) and from a logarithmic distribution between 0.1 and 1 for threshold parameters (K). Each parameter set was simulated for 100h using the ode15s integrator in Matlab and classified as mono-allelic, if Xist(Xi)>10*Xist(Xa) at the end of the simulation. For the cXR (*cis*-acting *Xist* repressor) model 32% of parameter sets maintained mono-allelic expression, but none of the other model was able to do so.

#### 1.3 Two-regulator models

To test whether other regulator classes than cXR would be able to maintain mono-allelic *Xist* expression, when combined with a second regulator, we built another 28 networks, each containing two different regulators A and B. Equation (1) in section 1.1 was thus modified as follows.

|  | Chr. X<sub>1</sub> | Chr. X<sub>2</sub> |  |
| --- | --- | --- | --- |
| **Xist** | $\frac{dx_1}{dt} = f(r_A) \cdot f(r_B) - x_1$ | $\frac{dx_2}{dt} = f(r_A) \cdot f(r_B) - x_2$ | (3) |

The resulting models have between 4-8 parameters depending on whether the regulators are silenced or escape.

#### 1.4 Simulating mono-allelic expression in two-regulators models

For each of the 28 two-regulator models >1,000 randomly chosen parameter sets were simulated as described in section 1.2. As shown in table M1 only networks containing a cXR can maintain mono-allelic expression.

**Table M1.** For each model containing either one regulator (diagonal) or two regulators the percentage of parameter sets that maintain mono-allelic expression are given.

|  | cXA | ecXA | tXA | etXA | cXR | ecXR | tXR | etXR |
| --- | --- | --- | --- | --- | --- | --- | --- | --- |
| cXA | 0 | 0 | 0 | 0 | 16.0 | 0 | 0 | 0 |
| ecXA |  | 0 | 0 | 0 | 30.1 | 0 | 0 | 0 |
| tXA |  |  | 0 | 0 | 26.0 | 0 | 0 | 0 |
| etXA |  |  |  | 0 | 29.9 | 0 | 0 | 0 |
| cXR |  |  |  |  | 32.1 | 2.7 | 6.3 | 2.1 |
| ecXR |  |  |  |  |  | 0 | 0 | 0 |
| tXR |  |  |  |  |  |  | 0 | 0 |
| etXR |  |  |  |  |  |  |  | 0 |

#### 1.5 Simulating male cells and bi-allelic expression in females

Since *Xist* is expressed from exactly one chromosome in each female cell, the underlying network should maintain mono-allelic expression, but destabilize the bi-allelic state in female cells and prevent *Xist* expression in male cells with only one X chromosome. To test which network would fulfill these criteria, three additional simulations were performed, where female cells initiated from an XiXi state (simulation 3), where male cells initiated from an Xi (simulation 4) or from an Xa state (simulation 2). To simulate male cells the models were modified as follows such that only a single X chromosome would be present.

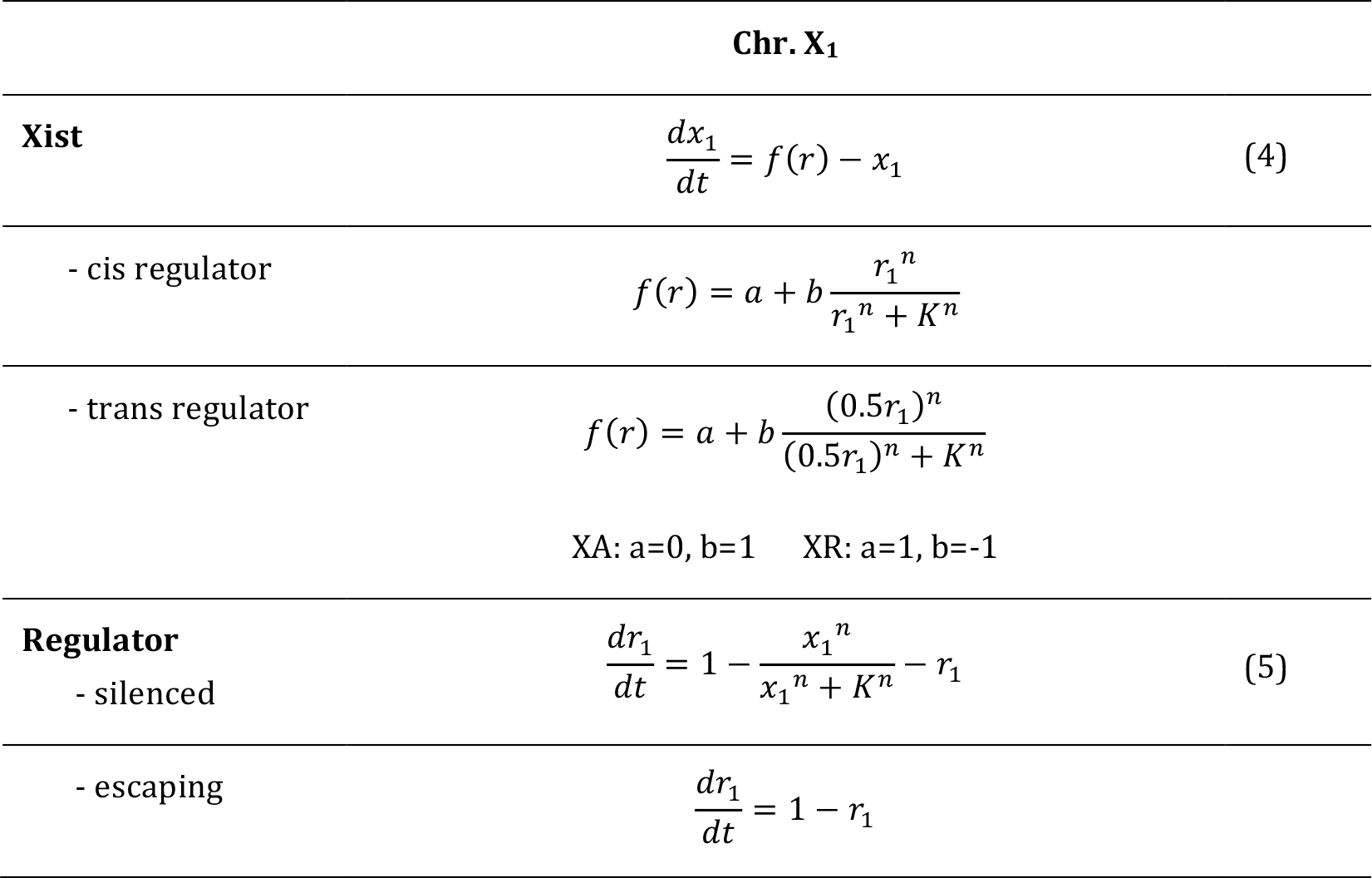

For each mono-allelic parameter set identified in sections 1.2 and 1.4 three additional simulations were performed with the following initial conditions.

|  | Model | Xist-1 | Xist-2 | Regulator-1 | Regulator-2 |
| --- | --- | --- | --- | --- | --- |
| **XiXi** | female | 0.99 | 1 | 0.01 (silenced)<br>1 (escaping) | 0.01 (silenced)<br>1 (escaping) |
| **Xi** | male | 1 |  | 0.01 (silenced)<br>1 (escaping) |  |
| **Xa** | male | 0.01 |  | 1 |  |

Based on the steady state of 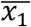 (Xist^high^ state) in the mono-allelic simulation in section 1.2, parameter sets were classified as follows.

| class | simulation | rule |
| --- | --- | --- |
| XiXi unstable | XiXi | $\bar{x}_1 < 10 \cdot Xist^{high}$ OR $\bar{x}_2 < 10 \cdot Xist^{high}$ |
| Xi unstable | Xi | $\bar{x}_1 < 10 \cdot Xist^{high}$ |
| Xa stable | Xa | $\bar{x}_1 < 10 \cdot Xist^{high}$ |

As shown in table M2 only a single model, namely the cXR-tXA model could maintain the XaXi and Xa states in female and male cells, respectively, while destabilizing both the XiXi and Xi states. All parameter sets that destabilized the XiXi state also destabilized the Xi state in male cells.

**Table M2.** Percentage of parameter sets in each class among all parameter set that can maintain the XaXi state.

|  | cXR-cXA | cXR-ecXA | cXR-tXA | cXR-etXA | cXR-ecXR | cXR | cXR-tXR | cXR-etXR |
| --- | --- | --- | --- | --- | --- | --- | --- | --- |
| **XiXi unstable** | 0 | 0 | 35.6 | 0 | 0 | 0 | 0 | 0 |
| **Xi unstable** | 0 | 0 | 61.5 | 28.5 | 0 | 0 | 0 | 0 |
| **Xa stable** | 100 | 100 | 100 | 100 | 100 | 100 | 82.8 | 83.3 |

### 2 The cXR-tXA model

The cXR-tXA model contains a *cis*-acting positive feedback loop mediated by a *cis*-acting repressor (cXR) and a *trans*-acting negative feedback, mediated by a *trans*-acting activator (tXA). The analysis in section 1 shows that the cXR-tXA model can explain maintenance of the correct *Xist* expression pattern (post-XCI state) for a subset of parameter sets. In the next step it was tested whether and under which conditions the model could also reproduce mono-allelic up-regulation of *Xist*, where a transition from the XaXa state (pre-XCI) to the XaXi state (post-XCI) occurs. Since a symmetry-breaking event, initiated by stochastic fluctuations, is required to transition from a symmetric XaXa state to an asymmetric XaXi state, *Xist* up-regulation was simulated in a stochastic manner.

#### 2.1 Stochastic simulation of the cXR-tXA model

Because absolute levels do not affect the qualitative results of the deterministic simulations in section 1 the equations were formulated in a way that the levels of *Xist* and its regulators would be scaled between 0 and 1. In stochastic simulations by contrast absolute abundances can strongly affect the allelic variability. To perform the simulations in a realistic regime, scaling factors were added to the model in section 1 (p_21_, p_22_, p_23_) such that the maximal levels of *Xist* and its regulators would vary between 50 and 500 molecules per allele, such that the ODE formulation of the model is modified as follows.

|  | Chr. X<sub>1</sub> | Chr. X<sub>2</sub> |
| --- | --- | --- |
| **Xist** | $\frac{dx_1}{dt} = p_{21} \cdot f(cXR_1) \cdot f(tXA) - x_1$ $f(cXR_1) = 1 - \frac{cXR_1^{p_{13}}}{cXR_1^{p_{13}} + (p_{22} \cdot p_{14})^{p_{13}}}$ | $\frac{dx_2}{dt} = p_{21} \cdot f(cXR_2) \cdot f(tXA) - x_2 \quad (6)$ $f(cXR_2) = 1 - \frac{cXR_2^{p_{13}}}{cXR_2^{p_{13}} + (p_{22} \cdot p_{14})^{p_{13}}}$ |
| | $f(tXA) = \frac{(0.5(tXA_1 + tXA_2))^{p_{11}}}{(0.5(tXA_1 + tXA_2))^{p_{11}} + (p_{23} \cdot p_{12})^{p_{11}}}$ | |
| **tXA** | $\frac{dtXA_1}{dt} = p_{23} \left[ 1 - \frac{x_1^{p_3}}{x_1^{p_3} + (p_{21} \cdot p_4)^{p_3}} \right] - tXA_1$ | $\frac{dtXA_2}{dt} = p_{23} \left[ 1 - \frac{x_2^{p_3}}{x_2^{p_3} + (p_{21} \cdot p_4)^{p_3}} \right] - tXA_2 \quad (7)$ |
| **cXR** | $\frac{dcXR_1}{dt} = p_{22} \left[ 1 - \frac{x_1^{p_5}}{x_1^{p_5} + (p_{21} \cdot p_6)^{p_5}} \right] - cXR_1$ | $\frac{dcXR_2}{dt} = p_{22} \left[ 1 - \frac{x_2^{p_5}}{x_2^{p_5} + (p_{21} \cdot p_6)^{p_5}} \right] - cXR_2 \quad (8)$ |

Similarly the kinetics of gene silencing do not effect the steady states of a system that were analyzed above in section 1, but will influence the ability to switch between states. Therefore two additional parameters were added to describe how long after *Xist* up-regulation cXR (p_7_) and tXA (p_8_) will be silenced. This is implemented by assuming that *Xist* will transition through p_7_/p_8_ silencing intermediates x’, x’’, x’’’… with a rate of 1h^−1^ before reaching the silencing competent states, x_1_/x_2_ (Fig. M1).

**Figure M1.**
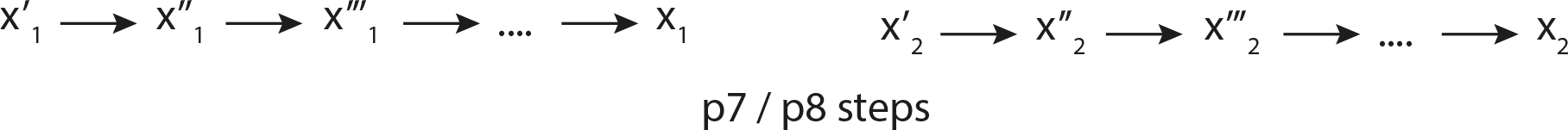
Schematic representation of the implementation of a silencing delay.

#### 2.2 Simulating mono-allelic *Xist* up-regulation

To facilitate the subsequent parameter analysis all hill coefficients were set to 3 (p_3_=p_5_=p_11_=p_13_=3) and the ODE simulations described in section 1 were repeated for 100,000 parameter sets. Each parameter set that maintained the XaXi and Xa states, while destabilizing the XiXi and Xi states (13.5%) was combined with 10 sets of silencing delays p_7_ and p_8_ chosen randomly between 1 and 20 and scaling factors p_21_, p_22_ and p_23_chosen randomly between 50 and 500. As shown in table M3, the resulting model has 9 parameters that were varied between simulations.

**Table M3.**
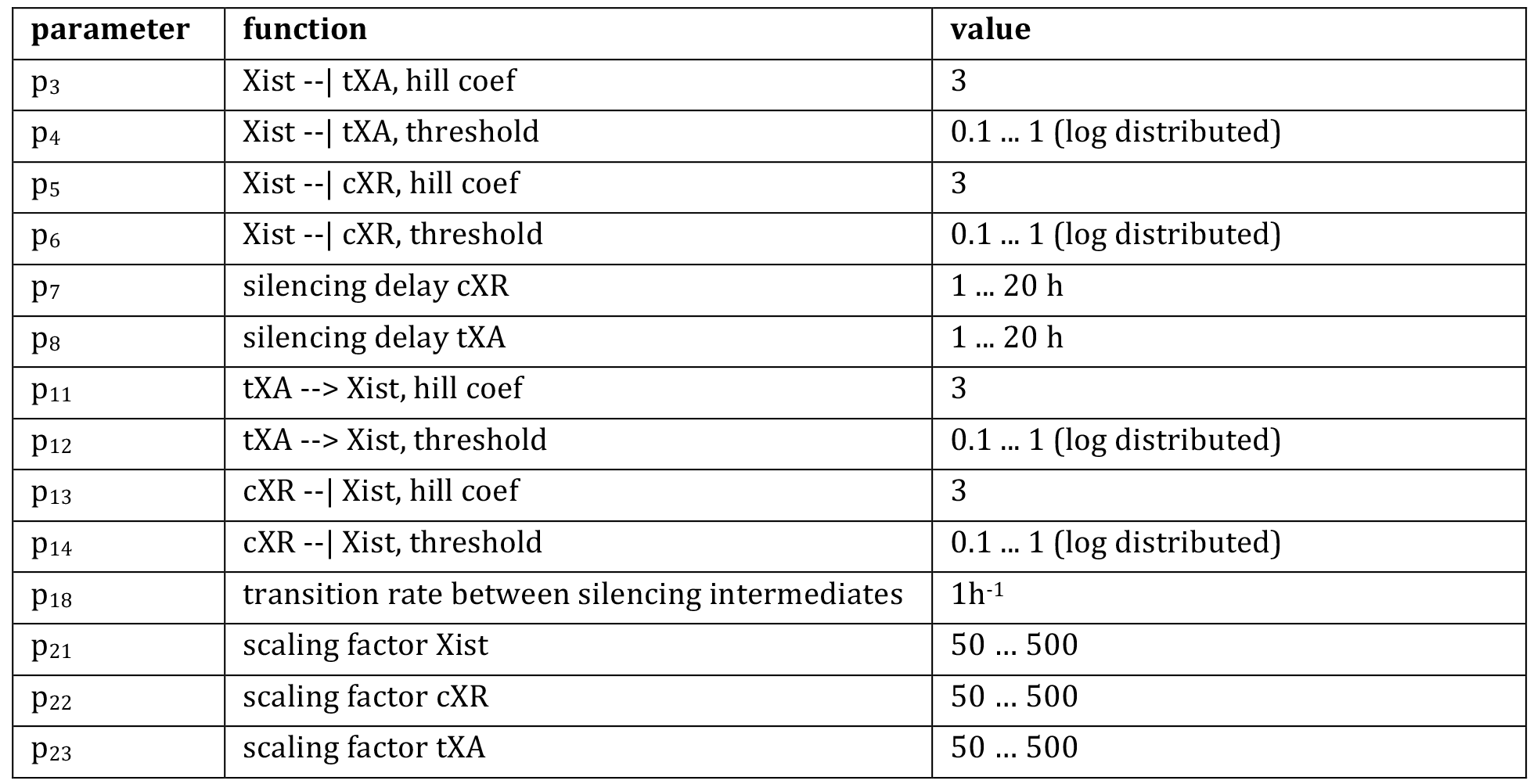
Parameters in the tXA-cXR model

For each parameter set, 100 cells were simulated for 100h starting from an XaXa state (x_1_=x_2_=0, cXR_1_=cXR_2_= p_22_, tXA_1_=tXA_2_= p_23_). The simulations were performed in Julia using the Gillespie algorithm^1^ and run on a computing cluster. Each chromosome was classified as *Xist* positive (Xist+) or negative (Xist−) for each hour of the simulation, depending on whether the mean *Xist* level exceeded 20% of the Xist^high^ state estimated from the ODE simulations above (0.2*Xist^high^*p_21_). Based on this classification the mean fraction of cells exhibiting mono- and bi-allelic expression was calculated. A small percentage of parameter sets tested (0.38%) could recapitulate efficient mono-allelic *Xist* up-regulation (>80% cells in the Xist+Xist− state during the last 20h of the simulation) as shown in Figure M2.

**Figure M2.**
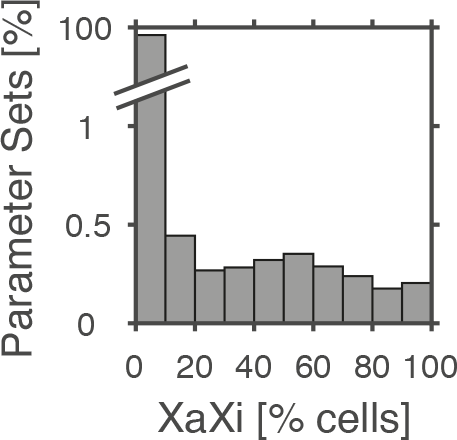
Distribution of the percentage of parameter sets that reproduce mono-allelic *Xist* up-regulation.

#### 2.3 Steady state analysis

To understand the prerequisites for mono-allelic *Xist* up-regulation in the cXR-tXA model we analyzed the steady states of the system, both locally at the allele level and globally at the cell level. The steady state levels of *Xist* for an individual allele were identified through an ODE simulation (see section 1) starting from different initial conditions (x=0, 0.1, 0.2…1, cXR=1, 0.9, 0.8,…0) and for different tXA doses, which remained constant during the simulation. The steady states reached after 100h of simulation are shown in Fig. 2E (top) in the main text for an example parameter set (p_4_=0.12, p_6_=0.15, p_12_=0.39, p_14_=0.40). tXA doses corresponding to 0, 1 or 2 active X-chromosomes (XiXi, XaXi, XaXa) are indicated, as calculated from the Xist^high^ and Xist^low^ steady state as follows.

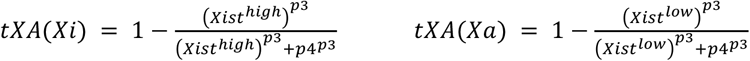

To identify the global steady states of an entire cell with two X chromosomes, the cXR-tXA ODE model was simulated for all combinations of initial values for x_1_=0, 0.1, 0.2,…1 and x_2_=_0_, 0.1, 0.2,…1. cXR_1_ and tXA_1_ were set to 1-x_1_ and cXR_2_ and tXA_2_ were set to _1_-x_2_. The steady states reached after 100h of simulation are shown for an example parameter set in Fig. 2E (bottom) in the main text. Steady states that can only be reached from a symmetric initial condition (x1=x2) are indicated as unstable (open circles).

To investigate the roles played by the two regulatory modules mediated by tXA and cXR, the effect on perturbing each module on system’s steady states was analyzed (Fig.2F-G in the main text). To perturb the positive feedback the threshold level of cXR required to repress *Xist* was set to a high value such that cXR is not any more involved in *Xist* regulation (p_14_=1000). To perturb the tXA modules, tXA was set to a constant value corresponding to a single dose present in the XaXi state as described above.

Surprisingly, we found a subset of parameter sets that could reproduce mono-allelic *Xist* up-regulation, where the system exhibited local and global bistability also in the presence of a double tXA dose. In these cases, the low steady state of *Xist* was rather unstable such that small fluctuations would allow a transition to the high state.

#### 2.4 Parameter rules

To understand, which parameter sets could reproduce mono-allelic *Xist* up-regulation, we analyzed the distribution of the parameter values among parameter sets that could maintain the correct *Xist* expression state in section 1 and that could simulate mono-allelic up-regulation in section 2.2 (Fig. M3).

**Fig. M3.**
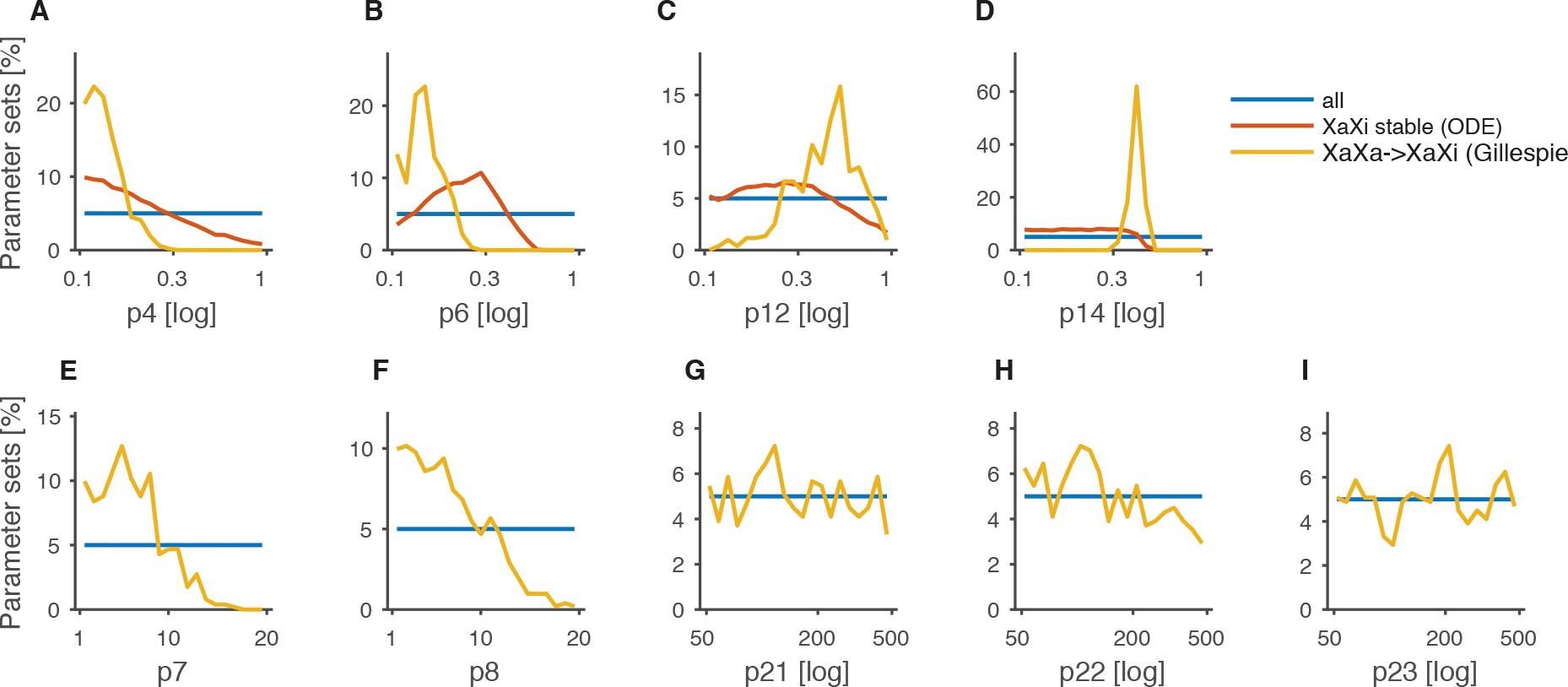
Distributions of parameter values across all tested parameter sets (blue), across all parameter sets that could maintain the XaXi state in the ODE simulation (red) and across all parameter sets that could reproduce mono-allelic *Xist* up-regulation. Silencing delays (p_7_ and p_8_) and scaling factors (p_21_, p_22_, p_23_) are only present in the stochastic model, but not in the ODE model.

The parameters p_4_ and p_6_ represent the silencing threshold for tXA and cXR, respectively. They are equal to the *Xist* level where tXA/cXR will be reduced to half of their maximal levels. Their values must be rather low to ensure efficient silencing upon mono-allelic *Xist* up-regulation (Fig. M3A+B). The parameters p_12_ and p_14_ describe how tXA and cXR, respectively, control *Xist*. They represent the tXA/cXR level where *Xist* activation/repression will be half-maximal. Both are required at intermediate levels for mono-allelic *Xist* up-regulation to allow *Xist* to respond sensitively to changes in tXA and cXR (Fig. M3C+D). The effect of p_12_ and p_14_ are discussed in more detail below. The silencing delays p_7_ and ps should be short (Fig. M3E+F), such that the Xist^high^ state can be locked in efficiently through cXR silencing and that the system can move the bi-stable regime through tXA silencing. For the scaling factors p_21_, p_22_ and p_23_ that determine the absolute levels of Xist, cXR and tXA, respectively, all values tested between 50 and 500 are compatible with Xist up-regulation (Fig. M3 G-I).

Both p_12_ and p_14_ must assume intermediate levels to allow *Xist* up-regulation (Fig. M3 C+D). In particular p_14_ must lie in a precise parameter range. To better understand these parameter requirements, we analyzed how these parameters affect the steady states of the system by visualizing the steady states in the Xist-cXR phase space. To this end, equations (6) and (8) were solved for steady state conditions 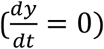 to

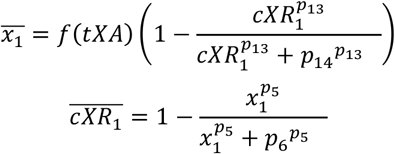

For one example parameter set that can recapitulate the XaXa-XaXi transition, we plot Xist (x1) vs cXR for both equations for 3 different doses of tXA, which would correspond to the XaXa (2x), XaXi (1x) and XiXi (0x) states (Fig. M4).

**Fig. M4.**
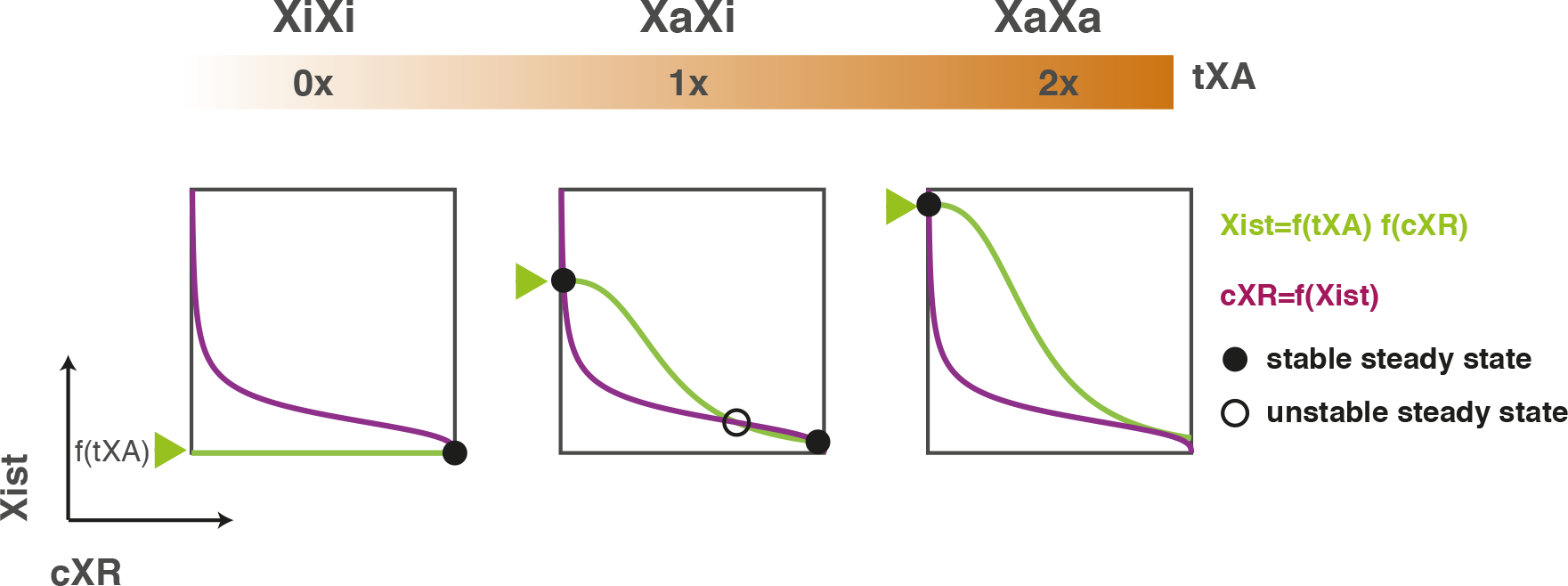
Xist-cXR phase space for one example parameter set for different tXA doses as indicated on top.

The two curves intersect at the system’s steady states. At the mono-allelic XaXi state (middle) the system exhibits bistability at the allele level. Because both a low and a high steady state exist for *Xist*, the two chromosomes can be stably maintained in two alternative states. In a putative XiXi state (left), where tXA is silenced on both chromosomes only the low state is stable. As a consequence *Xist* would be down-regulated if both chromosomes silence tXA, such that an XiXi state cannot be maintained in the system. Prior to the onset of X inactivation, both copies of tXA are active (XaXa) and only a single high steady state for *Xist* is present (right). This disappearance of the Xist^low^ state is the prerequisite for *Xist* up-regulation in the presence of a double tXA dose. In male cells that have only a single tXA dose (middle) the low state exists, thus preventing *Xist* up-regulation.

To understand how p_12_ and p_14_ would affect the system’s steady states, we plotted the phase diagram again for a two-fold increase or decrease in p_12_ or p_14_, respectively (Fig. M5)

**Fig. M5.**
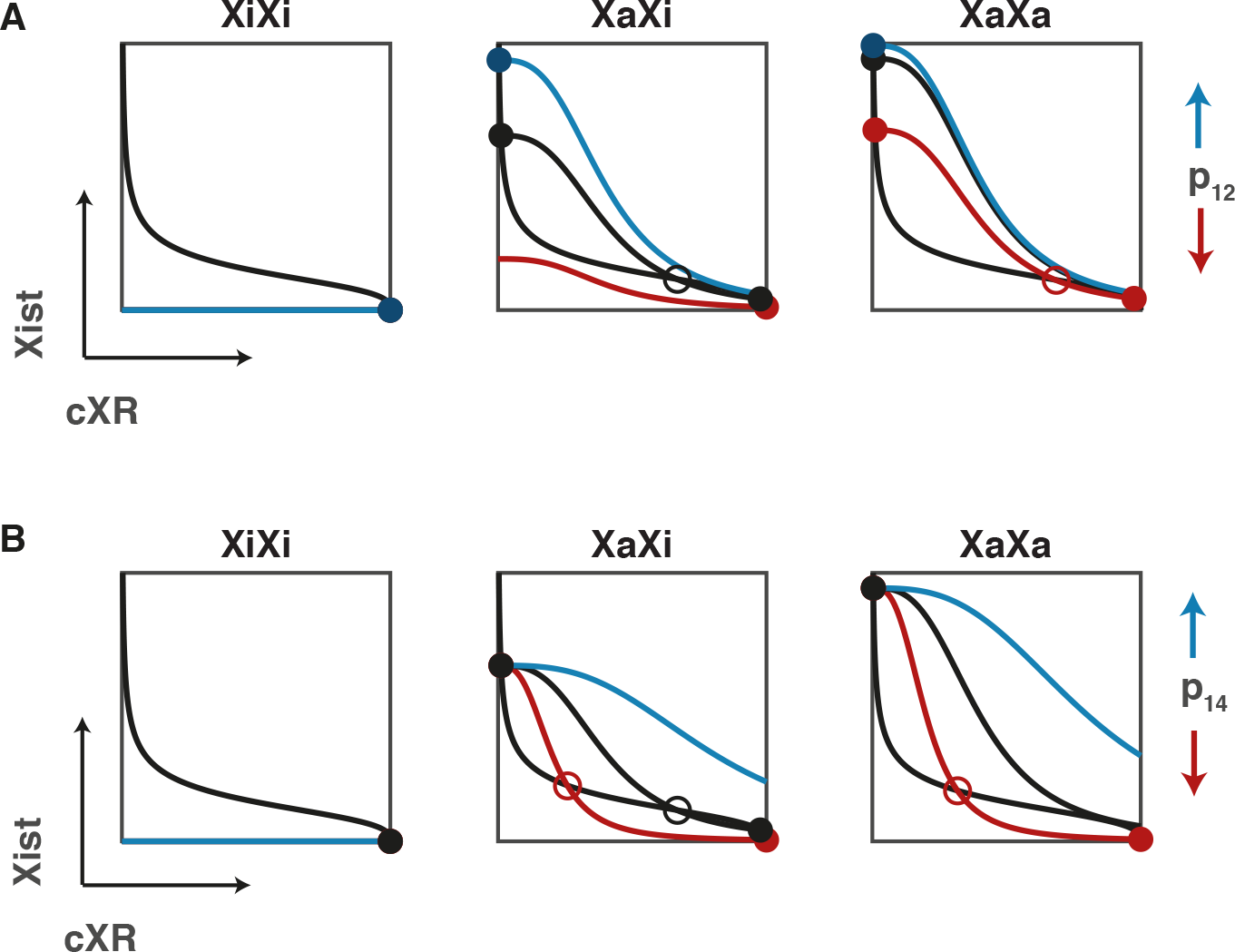
Xist-cXR phase space in response to changes in p_12_ (A) and p_14_ (B) for the same parameter set shown in Fig. M4. Blue indicates a two-fold increase of the respective parameter value and red a two-fold decrease.

An increase of p_12_ will result in disappearance of the Xist^low^ state in the XaXi scenario (Fig. M5A, middle, blue), while a decrease will result in loss of the Xist^high^ state (red). Therefore, p_12_ is required at intermediate levels to maintain and establish the mono- allelic state (cp. Fig. M3C). For p_14_, an increase will lead to loss of the Xist^low^ state (Fig. M5B, middle, blue). As a consequence, p_14_ must be below a certain level to allow bistability and thus maintenance of the XaXi state (cp. Fig. M3D, red). A decrease of p_14_by contrast results in the appearance of an Xist^low^ state in the XaXa scenario (Fig. M5B right, red), thus preventing *Xist* up-regulation in the pre-XCI state. Therefore, mono-allelic *Xist* up-regulation requires p_14_ to lie in a precise window (Fig. M3D).

#### 2.5 Simulating aneuploid and polyploid cells

To simulate cells that are mono-, tri- or tetrasomic for the X chromosome, *Xist* up-regulation was simulated as described in section 2.2 except that each cell contained one, three or four X chromosomes, each contributing a single tXA dose. For 50 randomly chosen parameter sets that were able to reproduce mono-allelic up-regulation in diploid cells, 100 cells were simulated. Each cell was classified as no Xi, Xi, XiXi, XiXiXi or XiXiXiXi depending on how many Xist+ alleles were present during the last 20 h of the simulation (Fig. 2H in the main text).

To simulate triploid (3n3X) and tetraploid cells (4n4X) we assumed either that tXA is repressed by autosomal transcription factors (Fig. 2I in the main text) or that an increase in nuclear size would result in an effective dilution of tXA compared to diploid cells (Fig. S1). To simulate autosomal tXA repression, equation (7) was substituted by

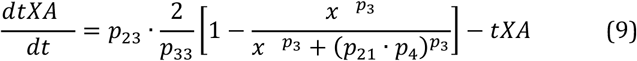

where p_33_ indicates the autosomal ploidy (3 for diploid cells, 4 for tetraploid cells). To simulate tXA dilution, equation (6) was substituted by

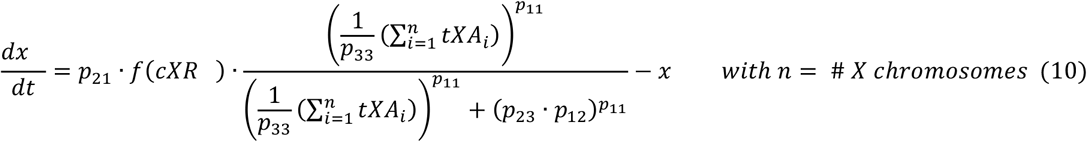

The four X chromosomes in a tetraploid cell and the three X chromosomes in a triploid cell would therefore produce the same effective tXA concentration as the two X’s in a diploid cell. The simulations were performed as described for polysomic cells above.

#### 2.6 Transient bi-allelic *Xist* up-regulation in mice, humans and rabbits

To investigate, whether the cXR-tXA model could reproduce transient bi-allelic *Xist* up-regulation as observed in rabbit embryos, we quantified the frequency of bi-allelic *Xist* expression (Xist+Xist+) across time for each parameter set that reproduced mono-allelic *Xist* up-regulation (in the last 20h of the simulation) in >80% cells in the simulations in section 2.2. The maximal level of bi-allelic expression observed was highly variable across parameter sets (Fig. M6).

**Fig. M6.**
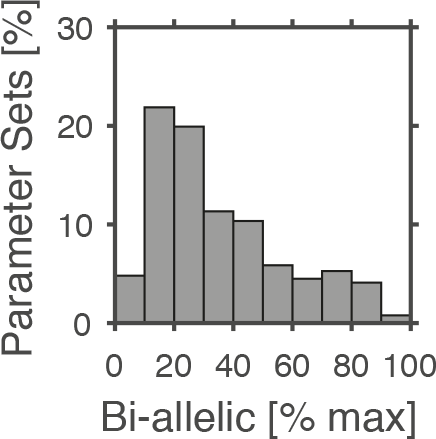
Distribution of the maximal percentage of cells that exhibit bi-allelic *Xist* expression during the simulation, for all parameter sets, where *Xist* is up-regulated mono-allelically in >80% of cells.

The level of transient bi-allelic *Xist* expression in the simulation depended on how fast *Xist* was up-regulated and how fast tXA was silenced. To quantify the speed of *Xist* up-regulation, we defined the “Switch-ON time” as the time point when one X chromosome had reached the Xist+ state (as defined in section 2.2). If switch-on did not occur before the end of the simulation it was set to the total simulation time (100h). The ratio of the average switch-ON time and the tXA silencing delay (p_8_) is inversely correlated with the maximal level of bi-allelic expression observed (Fig. 3A in the main text).

To understand whether cXR-tXA model could also explain extended bi-allelic *Xist* expression in human embryos, we tested whether and under which conditions absent or reduced silencing ability of *Xist* would lead to bi-allelic *Xist* expression. We identified two scenarios that could reproduce sustained bi-allelic expression: (1) cXR up-regulation/activation and (2) cXR dampening. To simulate these scenarios equations (7) and (8) were modified as follows and 100 parameter sets that could reproduce mono-allelic *Xist* expression were simulated as described in section 2.2.

|  | (1) cXR upregulation/activation | (2) cXR dampening |
| --- | --- | --- |
| **tXA** | $\text{for } t \leq t1:$ $\frac{dtXA_i}{dt} = p_{23} - tXA_i$<br>$\text{for } t > t1:$ $\frac{dtXA_i}{dt} = p_{23} \left[ 1 - \frac{x_i^{p_3}}{x_i^{p_3} + (p_{21} \cdot p_4)^{p_3}} \right] - tXA_i$ | $\text{for } t \leq t1:$ $\frac{dtXA_i}{dt} = p_{23} - tXA_i$<br>$\text{for } t > t1:$ $\frac{dtXA_i}{dt} = p_{23} \left[ 1 - \frac{x_i^{p_3}}{x_i^{p_3} + (p_{21} \cdot p_4)^{p_3}} \right] - tXA_i$ |
| **cXR** | $\text{for } t < t1:$ $\frac{dcXR_i}{dt} = 0 - cXR_i$<br>$\text{for } t \geq t1:$ $\frac{dcXR_i}{dt} = p_{22} \left[ 1 - \frac{x_i^{p_5}}{x_i^{p_5} + (p_{21} \cdot p_6)^{p_5}} \right] - cXR_i$ | $\text{for } t < t1$ $\frac{dcXR_i}{dt} = p_{22} \left[ 1 - p_{34} * \frac{x_i^{p_5}}{x_i^{p_5} + (p_{21} \cdot p_6)^{p_5}} \right] - cXR_i$<br>$\text{for } t \geq t1:$ $\frac{dcXR_i}{dt} = p_{22} \left[ 1 - \frac{x_i^{p_5}}{x_i^{p_5} + (p_{21} \cdot p_6)^{p_5}} \right] - cXR_i$ |

In scenario 1 it was assumed that cXR was not expressed before t1 (=day 4) and that tXA could not be silenced by *Xist* during that period. In scenario 2, we assumed that cXR is subject to dampening by *Xist* RNA while tXA is unaffected by *Xist.* To simulate dampening of cXR by *Xist* we introduced an additional parameter p_34_ that describes maximal reduction of cXR by *Xist*. For each parameter set, p_34_ was randomly drawn from a uniform distribution between 0.01 and 0.99. Both scenarios could reproduce extended bi-allelic expression and the transition to the mono-allelic state (Fig. 3D in the main text and supplemental Fig. S1).

#### 2.7 Simulating experimental modulation of bi-allelic *Xist* up-regulation

To simulate induction of *Xist* with doxycycline equation (6) was modified by introducing two additional parameters per chromosome as follows.

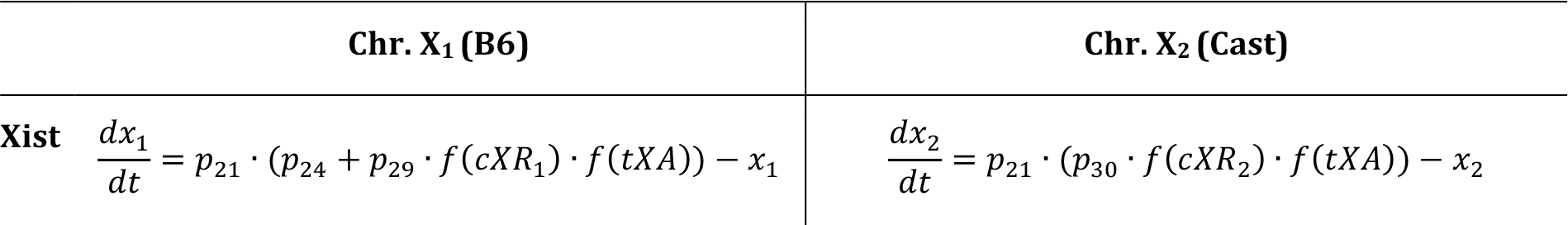

|  | Chr. X<sub>1</sub> (B6) | Chr. X<sub>2</sub> (Cast) |
| --- | --- | --- |
| **Xist** | $\frac{dx_1}{dt} = p_{21} \cdot (p_{24} + p_{29} \cdot f(cXR_1) \cdot f(tXA)) - x_1$ | $\frac{dx_2}{dt} = p_{21} \cdot (p_{30} \cdot f(cXR_2) \cdot f(tXA)) - x_2$ |

P_24_ describes *Xist* regulation by doxycycline on the B6 chromosome and is set to 10 if Doxycycline is present or to 0 if doxycycline is absent. P_29_ and p_30_ control regulation of *Xist* by tXA and cXR and are set to 1 on an allele that is not induced with doxycycline but to 0 if induced with doxycycline, since *Xist* expression then becomes independent of regulation by tXA and cXR. Before the start of differentiation p_29_ and p_30_ are also set to 0 representing the action of stem cell specific factors that prevent upregulation of *Xist* in undifferentiated cells by repressing *Xist*. Parameter settings to simulate doxycycline treatment are shown below.

|  | + Dox |  | Control |  |
| --- | --- | --- | --- | --- |
| | $t < 0$ | $t \geq 0$ | $t < 0$ | $t \geq 0$ |
| p<sub>24</sub> | 10 | 10 | 0 | 0 |
| p<sub>29</sub> | 0 | 0 | 0 | 1 |
| p<sub>30</sub> | 0 | 1 | 0 | 1 |

To simulate skewing in the TX1072dT line due to a heterozygous Tsix deletion on the Cast chromosome p_29_ was set to 33% of p_29_, a summary of the parameter setting is shown below.

| | $t < 48h$ | $t \geq 48h$ |
| --- | --- | --- |
| p<sub>24</sub> | 0 | 10 |
| p<sub>29</sub> | 0.33 | 0 |
| p<sub>30</sub> | 1 | 1 |

### 3 Model with *Tsix*-mediated local positive feedback

#### 3.1 Model description

Reactions describing transcription initiation, transcription elongation and RNA degradation of *Xist* and *Tsix* were combined into a mathematical model (Fig. M7).

**Fig. M7.**
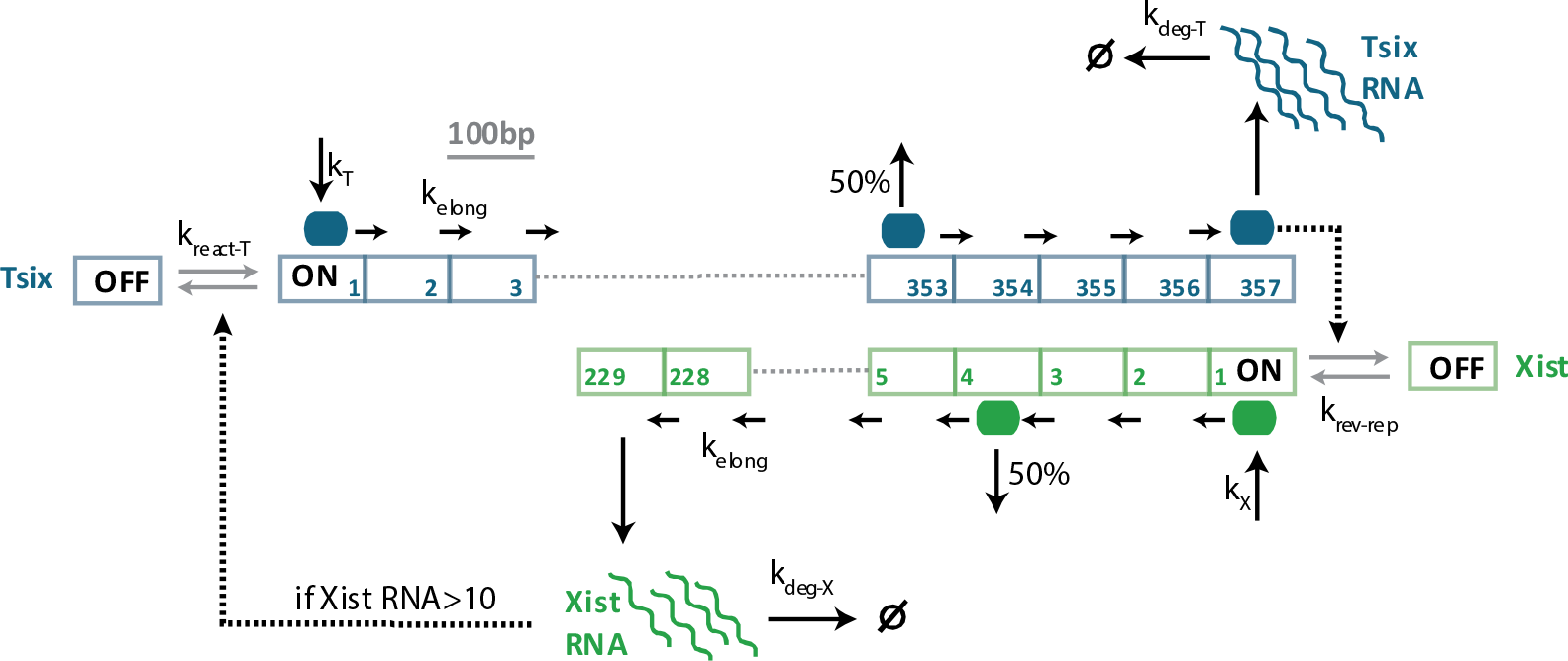
Schematic representation of antisense simulation.

Both promoters were assumed to exist in an ‘OFF’ state, where no transcription occurs, and an, ON’ state, where transcription is initiated with constant transcription rates k_X_ and k_T_. The *Tsix* promoter is turned off by *Xist* RNA-mediated silencing, the *Xist* promoter is turned off by passing *Tsix* polymerases. The OFF state is reverted back to the ON state with rate k_rev-rep_. To describe transcriptional elongation, the *Xist* and *Tsix* gene bodies were divided into segments of 100nt and RNA polymerases move along the gene body with a constant rate (k_elong_). Fully elongated transcripts produce one RNA molecule. Degradation of the RNA obeys first-order reaction kinetics with the rates k_deg-X_ and k_deg-T_.

Transcription of *Xist* and *Tsix* mutually interfere by the following mechanisms: (i) transcriptional interference occurs between sense- and antisense transcribing polymerases with one randomly chosen polymerase being removed from the gene; (ii) *Tsix* polymerases transcribing through the *Xist* promoter region induce a transition to the OFF state of the *Xist* promoter that can be reverted to the ON state with k_rev-rep_ (the half live of the repressed state is given by 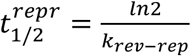); (iii) Silencing of the *Tsix* promoter by *Xist* RNA: If *Xist* RNA is present above a threshold level of 10 RNA molecules it induces a transition of the *Tsix* promoter to the OFF state.

To account for regulation of Xist by tXA, the tXA dosage was assumed to modulate the effective *Xist* initiation rate 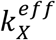 as follows:

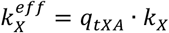

where k_X_ is the *Xist* initiation rate in the presence of a single tXA dose and q_tXA_=0,1,2 depending on whether no, one or two tXA loci are active. The tXA concentration was modeled as a step function with the value 1 if the respective tXA allele is active and the value 0 if the respective tXA allele has been silenced by *Xist* RNA. The kinetics of RNA and protein decay of tXA were not accounted for explicitly but were instead assumed to modulate the tXA silencing kinetics (see below).

To reproduce coupling of XCI to development, 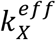 must be influenced by the differentiation timing, representing the action of stem cell specific factors that prevent *Xist* upregulation in undifferentiated cells by repressing *Xist*. The differentiation dependency was formulated as a step function that changes its value at the point of induction of differentiation such that 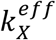 prior to differentiation was 10% of the k_X_ value after the onset of differentiation.

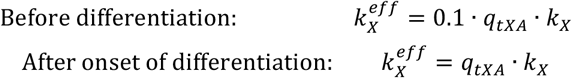

Since Xist-dependent silencing is known to occur with a delay of hours or days after *Xist* has been up-regulated^2,3^, we implemented a silencing delay described by the parameters sil_Tsix_ or sil_tXA_. To this end, each chromosome passes stochastically through a number of intermediate states S1, S2…Sn once *Xist* expression from that chromosome has exceeded a certain threshold (10 molecules) and gene silencing occurs once the final silencing state Sn has been reached. If the level of *Xist* RNA molecules drops below the threshold before Sn has been reached, the chromosome immediately passes back to the unsilenced state S0. The transitions through the intermediate states occur with rate 1h^−1^ such that the number of intermediate states given by the model parameters sil_Tsix_ or sil_tXA_ is equal to the mean silencing delay. Silencing is assumed to be reversed, if the *Xist* level drops below the threshold of 10 molecules. Reactivation of *Tsix* and tXA will then occur with a single stochastic reaction with the rates k_react-T_ and k_react-tXA_ respectively (see figure below).

**Fig. M8.**
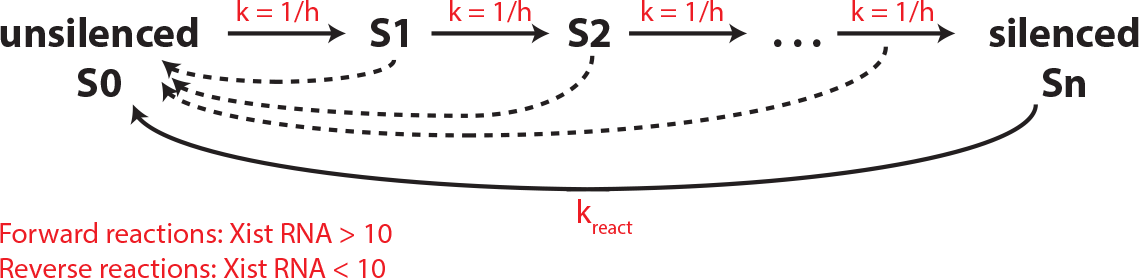
Schematic representation silencing and reactivation in the antisense model.

#### 3.2 Simulating maintenance of the XaXi state

To find parameter values that could reproduce mono-allelic *Xist* up-regulation, we first scanned a large parameter space for sets that would maintain the XaXi state. Subsequently we tested, whether those sets can reproduce the transition from the XaXa to the XaXi state. Degradation and elongation rates were set to fixed values based on previous experimental estimates (Table M4). For the *Xist* half-life previous studies have attempted an experimental estimation, resulting in values of 2 and 6h respectively^4,5^. We therefore use the mean (4h), which results in a degradation rate of 0.1733 h^−1^ (ln(2)/t_1/2_). Since silencing has already occurred in the XaXi state and silencing kinetics should therefore not affect the outcome of the simulation, silencing and reactivation rates were set to constant values (1h^−1^). Similarly, the tXA is present at a constant single dose in post-XCI cells with a single Xa and is therefore set to a constant value of 1. All other parameters were varied within realistic parameter ranges and systematically combined resulting in 8000 parameter sets in total (Table M4).

**Table M4.**
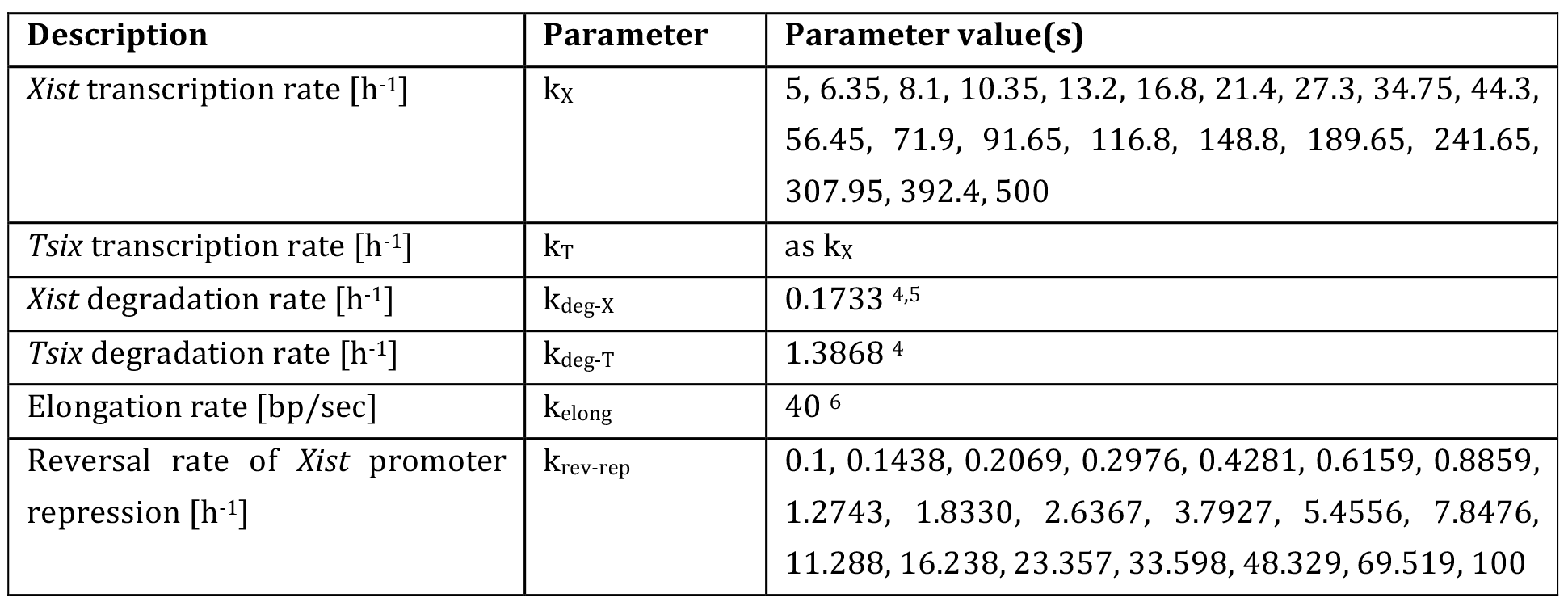
Parameter values in the antisense model

| Description | Parameter | Parameter value(s) |
| --- | --- | --- |
| *Xist* transcription rate [$\text{h}^{-1}$] | $k_X$ | 5, 6.35, 8.1, 10.35, 13.2, 16.8, 21.4, 27.3, 34.75, 44.3, 56.45, 71.9, 91.65, 116.8, 148.8, 189.65, 241.65, 307.95, 392.4, 500 |
| *Tsix* transcription rate [$\text{h}^{-1}$] | $k_T$ | as $k_X$ |
| *Xist* degradation rate [$\text{h}^{-1}$] | $k_{\text{deg-X}}$ | 0.1733 <sup>4,5</sup> |
| *Tsix* degradation rate [$\text{h}^{-1}$] | $k_{\text{deg-T}}$ | 1.3868 <sup>4</sup> |
| Elongation rate [bp/sec] | $k_{\text{elong}}$ | 40 <sup>6</sup> |
| Reversal rate of *Xist* promoter repression [$\text{h}^{-1}$] | $k_{\text{rev-rep}}$ | 0.1, 0.1438, 0.2069, 0.2976, 0.4281, 0.6159, 0.8859, 1.2743, 1.8330, 2.6367, 3.7927, 5.4556, 7.8476, 11.288, 16.238, 23.357, 33.598, 48.329, 69.519, 100 |

To investigate the stability of the Xa and Xi, each state was simulated using the initial conditions given in the table below. The RNA levels for transcribed genes was set to their maximal steady state value and the polymerase complexes were randomly distributed along the gene body.

|  | Xi | Xa |
| --- | --- | --- |
| *Xist* RNA | $k_X/k_{\text{deg-X}}$ | 0 |
| *Tsix* RNA | 0 | $k_T/k_{\text{deg-T}}$ |
| *Xist* promoter | ON | OFF |
| *Tsix* promoter | OFF | ON |
| # *Xist* polymerases | $\frac{L}{k_{\text{elong}}} * k_X$ (L=22 900 bp) | 0 |
| # *Tsix* polymerases | 0 | $\frac{L}{k_{\text{elong}}} * k_T$ (L=35 700 bp) |

In the simulation, transcription elongation occurs at fixed time intervals of 2.5 seconds inferred from measurements of polymerase speed (elongation of one 100bp interval at k_elong_=40bp/sec). Between elongation steps, all other reactions are simulated using the stochastic Gillespie algorithm^1^. For each parameter set 100 Xi/Xa pairs were simulated for 500h to reach the steady state.

A simulation was classified as stably maintaining the XaXi state if *Xist* was on average present with >10 molecules at the Xi and with <10 molecules at the Xa during the last 50h of the simulation. Parameter sets, where >99% of Xa/Xi pairs stably maintained the XaXi state were classified as mono-allelic.

#### 3.3 Simulating mono-allelic *Xist* up-regulation

To simulate the onset of X inactivation, both chromosome were initiated from the Xa state (see 3.2) in undifferentiated cells with double tXA dosage present (q_tXA_=2). *Xist* up-regulation was simulated for all parameter sets that could stably maintain the XaXi state in section the previous section (4001 sets). Each parameter set was combined with 500 combinations of randomly sampled values for sil_tXA_, sil_Tsix_, k_react-tXA_ and k_react-T_ (see following table).

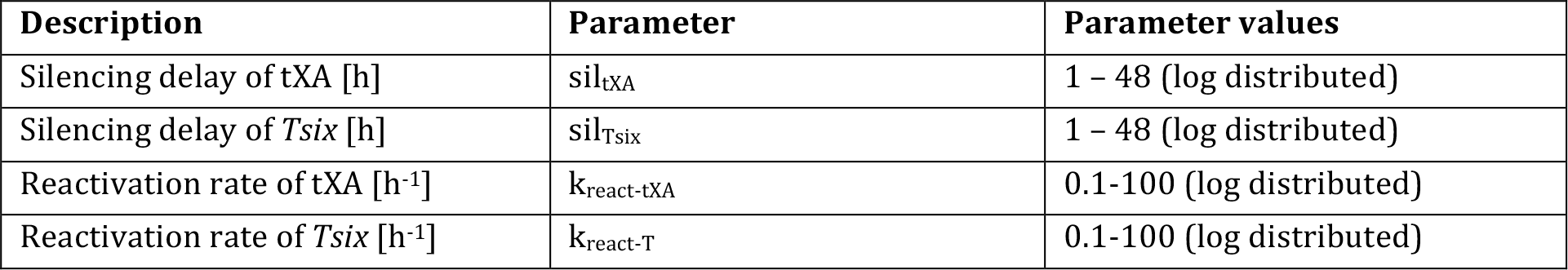

For each parameter set 100 cells were simulated. To reach the steady state prior to differentiation, the cells were simulated for 10h in an undifferentiated state, then 100 hours of differentiation were simulated as this is the relevant time scale of XCI. Each cell was classified as mono-allelic, if during the last 20h of the simulation >10 molecules of *Xist* RNA were present on average at one chromosome (Xi) and <10 molecules on the other (Xa). 1.17% of the tested parameter sets upregulated *Xist* mono-allelically in >99% cells and were thus classified as mono-allelic (XaXa->XaXi in Fig. 5D in the main text).

#### 3.4 Model simplification

To analyze which of the repressive mechanisms are required for establishment and maintenance of the Xa/Xi state, we systematically investigated reduced model structures with all three, a combination of two or only a single repressive mechanism. These simplifications were implemented as follows.

- In all models without *Xist* promoter repression, passing *Tsix* polymerases do not affect the *Xist* promoter state (parameter k_rev-rep_ removed).
- In all models without *Tsix* promoter silencing, the *Xist* RNA does not affect the *Tsix* promoter state.
- In all models without transcriptional interference (TI), *Xist* and *Tsix* polymerases were assumed to be able to bypass each other.

First, maintenance of the XaXi state was assessed for all six reduced models as described in section 3.2. For all parameter sets that could maintain the XaXi state, *Xist* up-regulation was simulated as described in section 3.3. A summary of the simulations of the full model and of all reduced models is given in table M5. Only the reduced model, where mutual inhibition of *Xist* and *Tsix* was mediated by silencing of the *Tsix* promoter and by TI could reproduce mono-allelic *Xist* up-regulation. This model was termed the “antisense model” and used for all subsequent simulations.

**Table M5.**
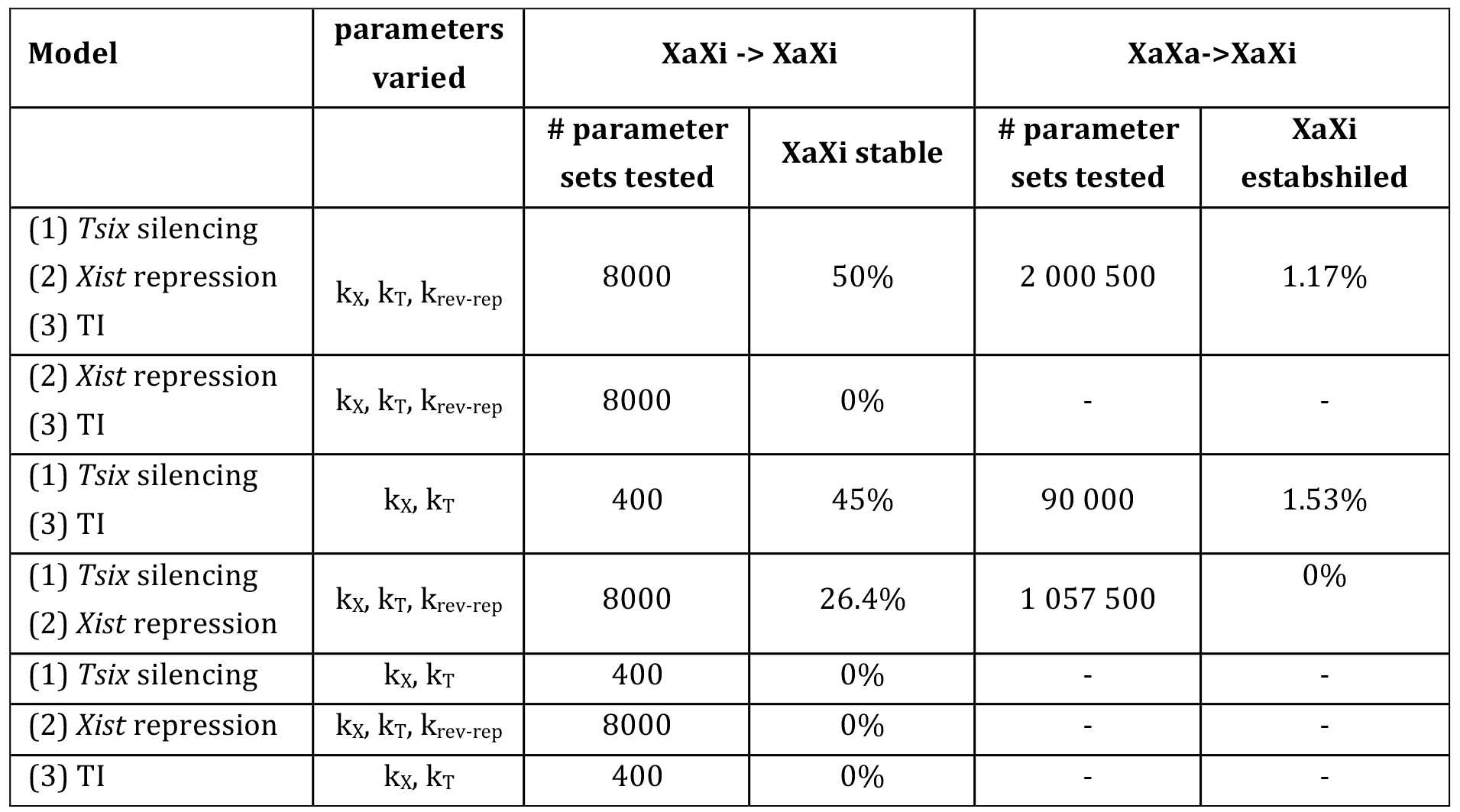
Summary of model reduction

#### 3.5 Simulations of *Xist* and *Tsix* mutant cell lines

To simulate experimental data we selected a subset of parameter sets from the simulation in 3.4 that robustly led to monoallelic *Xist* upregulation (>99% cells) and were in agreement with experimental observations according to the following constraints:

i. The maximal percentage of bi-allelically expressing cells over the simulated time course should be below 20%
ii. The mean *Xist* expression level must lie between 200 and 600 RNA molecules^4^.
iii. All cells up-regulate *Xist* within 48h after induction of differentiation.

Since only 34 parameter sets with unique k_X_, k_T_, sil_tXA_ and sil_Tsix_ combinations fulfilled these criteria, we performed another simulation to identify more parameter sets that could potentially simulate experimental data. To this end, the simulation in 3.4 was repeated with additional, randomly sampled values for k_X_ and k_T_. The parameter ranges for kX were set such that the steady state expression level of *Xist* (k_X_/k_X-deg_) was between 200 and 600 molecules. Since an analysis of the parameter sets identified in section 3.4 had revealed that mono-allelic *Xist* up-regulation requires a k_X_-to-k_T_ ratio between 0.4 and 0.8, k_T_ was sampled within this range. A total of 500,000 parameter sets were simulated. All parameters were sampled randomly within the ranges given in the following table.

| Description | Parameter | Parameter values |
| --- | --- | --- |
| *Xist* initiation rate [$\text{h}^{-1}$] | $k_X$ | 34-104 (log distributed) |
| *Tsix* initiation rate [$\text{h}^{-1}$] | $k_T$ | $k_X/0.8$-$k_X/0.4$ (lin distributed) |
| Silencing delay of tXA [h] | $\text{sil}_{\text{tXA}}$ | 0 – 48 (log distributed) |
| Silencing delay of *Tsix* [h] | $\text{sil}_{\text{Tsix}}$ | 0 – 48 (log distributed) |
| Reactivation rate of tXA [$\text{h}^{-1}$] | $k_{\text{react-tXA}}$ | 0.1-100 (log distributed) |
| Reactivation rate of *Tsix* [$\text{h}^{-1}$] | $k_{\text{react-T}}$ | 0.1-100 (log distributed) |

From these simulations 100 sets fulfilling the above requirements were randomly selected and were used in the following simulations. *Xist* and *Tsix* mutations were simulated as described above with the following modifications.

i. *Tsix*^+/−^: *Tsix* initiation rate k_T_=0 on one allele
ii. *Tsix*^−/−^: *Tsix* initiation rate k_T_=0 on both alleles.
iii. *Xist*^+/−^: *Xist* initiation rate k_X_=0 on one allele

For each of the 100 selected parameter sets 100 cells were simulated for each mutant. Fig. 7B-D in the main text shows the trajectory of one representative cell (top), the final expression state of all cells for one parameter set (middle) and the distributions of expression patterns observed across all tested parameter sets (bottom). To estimate the halftime of *Xist* up-regulation T_1/2_ shown in Fig. 7E (main text) we determined the earliest time point where 50% of simulated cells had up-regulated *Xist* (>10 molecules).

